# Neurodevelopmental defects in Dravet syndrome *Scn1a*^+/-^ mice: targeting GABA-switch rescues behavioral dysfunctions but not seizures and mortality

**DOI:** 10.1101/2024.03.06.583652

**Authors:** Lara Pizzamiglio, Fabrizio Capitano, Evgeniia Rusina, Giuliana Fossati, Elisabetta Menna, Isabelle Léna, Flavia Antonucci, Massimo Mantegazza

## Abstract

Dravet syndrome (DS) is a developmental and epileptic encephalopathy (DEE) caused by mutations of the *SCN1A* gene (Na_V_1.1 sodium channel) and characterized by seizures, motor disabilities and cognitive/behavioral deficits, including autistic traits. The relative role of seizures and neurodevelopmental defects in disease progression, as well as the role of the mutation in inducing early neurodevelopmental defects before symptoms’ onset, are not clear yet. A delayed switch of GABAergic transmission from excitatory to inhibitory (GABA-switch) was reported in models of DS, but its effects on the phenotype have not been investigated.

Using a multi-scale approach, here we show that targeting GABA-switch with the drugs KU55933 (KU) or bumetanide (which upregulate KCC2 or inhibits NKCC1 chloride transporters, respectively) rescues social interaction deficits and reduces hyperactivity observed in P21 *Scn1a^+/-^* DS mouse model. Bumetanide also improves spatial working memory defects. Importantly, neither KU nor bumetanide have effect on seizures or mortality rate. Also, we disclose early behavioral defects and delayed neurodevelopmental milestones well before seizure onset, at the beginning of Na_V_1.1 expression.

We thus reveal that neurodevelopmental components in DS, in particular GABA switch, selectively underlie some cognitive/behavioral defects, but not seizures. Our work provides further evidence that seizures and neuropsychiatric dysfunctions in DEEs can be uncoupled and can have differential pathological mechanisms. They could be treated separately with targeted pharmacological strategies.

## Introduction

Developmental and epileptic encephalopathies (DEEs) are characterized by severe epileptic seizures with early-onset and a variable amount of underlying neurodevelopmental impairment that tends to worsen as a consequence of epilepsy, but it is difficult to discriminate the neurodevelopmental and epileptic components of the disease ^1^. Dravet syndrome (DS) is a severe DEE ^2,3^ caused by heterozygous mutations of the *SCN1A* gene, which lead to loss of function of the Na_V_1.1 voltage-gated sodium channel ^4^. Disease onset occurs during the first year of life with seizures triggered by fever or hyperthermia, which persist also in adulthood, and subsequent appearance of various other types of seizures with high rate of sudden unexpected death in epilepsy (SUDEP) ^3,5^. Beyond the severe epileptic phenotype, DS is characterized by developmental plateauing with motor disabilities, cognitive deficits and neuropsychiatric symptoms that often overlap with symptoms of autism spectrum disorder (ASD), which are permanent and greatly contribute to the burden of the disease ^6–9,3^. Three stages have been proposed in the natural history of DS: febrile, worsening and stabilization ^5^. The available drugs are symptomatic treatments with limited efficacy and insufficient control of convulsive seizures, whereas there is not yet a treatment reported as clearly effective on cognitive/behavioral dysfunctions ^3,10^. Heterozygous knock-out Na_V_1.1 (*Scn1a*^+/-^) mice recapitulate the human DS phenotype ^11–13^. Hyperthermia-induced seizures can be induced from around postnatal-day 15 (P15), equivalent to the febrile stage. Onset of spontaneous recurrent seizures is at around P20 with high SUDEP rate, as well as cognitive/behavioral defects, including autistic-like traits; in this period, which is equivalent to the human worsening stage, seizures are particularly frequent and mortality particularly high. A stabilization of the phenotype is observed after around P35, but cognitive/behavioral defects are permanent, as in patients. Mouse models have been instrumental for showing that Na_V_1.1 loss-of-function, caused by DS mutations, induces reduction of sodium current and of excitability in GABAergic interneurons, leading to impaired GABAergic synaptic transmission ^12,14^. This mechanism has been confirmed by numerous studies in different animal models and in patients, showing the pivotal role of GABAergic interneurons’ dysfunctions in DS ^11,15–18^.

The relative contribution of seizures and underlying neurodevelopmental impairments to the encephalopathy, in particular to neuropsychiatric features, is still debated in DS, as for other DEEs. In fact, some clinical studies highlighted mainly epileptic encephalopathy features, whereas others showed developmental alterations even before the onset of epilepsy and reported that in some cases the severity of epilepsy and cognitive defects were not correlated ^19–23^.

Although hypoexcitability of GABAergic neurons is the most convincing pathological mechanism and therapeutic target in DS, the detailed molecular mechanisms underlying epilepsy and comorbidities, their causal relationships and the precise time window at which the different pathological features develop are still poorly understood, limiting the development of targeted therapeutic strategies. A study has shown that in a DS *Scn1a* knock-out mouse model the reversal potential of GABA-evoked currents is depolarized compared to WT mice between P13 and P21, consistent with a delay in the switch of GABAergic transmission from excitatory to inhibitory (i.e. GABA-switch) ^24^. Also, a depolarized reversal potential for GABA-evoked currents was observed in *Xenopus* oocytes transplanted with cell membranes prepared from autopsy brain tissues of DS patients ^25^. However, the effects of this dysfunction on the DS phenotype were not investigated, although delayed mortality rescuing GABA-switch was reported in a model of sodium channel β1 subunit DEE ^24^, which is a different clinical entity with different phenotype and pathological mechanisms compared to DS ^1,26,27^.

The GABA-switch is a fundamental step of brain development that reflects the maturation time course of GABAergic signaling; it is in general completed by P15 and depends on the electrochemical Cl^-^ gradient, which is set by the relative expression of cation–chloride transporters NKCC1 (Na^+^-K^+^-Cl^-^ cotransporter isoform 1) and KCC2 (K^+^-Cl^-^ cotransporter isoform 2)^28–30^. Importantly, a delay in GABA-switch has been found in numerous models of neurodevelopmental disorders and it has been proposed as a therapeutic target in patients ^31–33^. Overall brain expression of Na_V_1.1 starts around P8 in mice and homozygous *Scn1a*^-/-^ mice die by P15 ^12,16^, showing that Na_V_1.1 loss of function can induce early effects. Thus, GABA-switch may be a target in therapeutic approaches for DS, and early neurodevelopmental defects may be implicated in DS.

Here, we investigated in DS *Scn1a*^+/-^ mice overall neurodevelopmental defects in the period before the onset of seizures, and we targeted GABA-switch in DS *Scn1a*^+/-^ mice by performing specific pharmacological interventions and assessing their effect on both epileptic and behavioral phenotypes.

## Methods

### Animals

Mice were group-housed in standard mouse cages, on a 12-h day/12-h night cycle and allowed free access to food and water. We used the exon 25-floxed (Fl) *Scn1a* mice (purchased from MMRRC, Stock No: 041829-UCD) ^15,34^, which was maintained on a C57BL/6J background. They were genotyped between post-natal day 5 (P5) and P7 for the *Scn1a* Fl allele using the primers 5′-CTTGATGTGTTGAAATTCAC-3′ and 5′-TATAGAGTGTTTAATCTCAAC-3′. Meox2-Cre mice were purchased from the Jackson Laboratory (Stock No: 003755) and maintained on a C57BL/6J background. They were genotyped at P5-P7 for the presence of Cre using the primers 5′-GGTTTCCCGCAGAACCTGAA-3′ and 5′-CCATCGCTCGACCAGTTTAGT-3′. In the Meox2-Cre line, Cre recombinase is expressed in all the tissues: it is observed in the epiblast as early as embryonic day 5 and subsequently in all the derivatives of the epiblast (Endoderm, Mesoderm and Ectoderm). For the experiments, we used F1 mice generated mating Fl *Scn1a* homozygous females and Meox2-Cre heterozygous males, comparing *Scn1a*^Fl/+^:Meox2-Cre^+/-^ mice as DS model (*Scn1a*^+/-^) and *Scn1a*^Fl/+^:Meox2-Cre^-/-^ mice as wild-type controls (WT). This strategy has been already used in previous studies and *Scn1a*^Fl/+^:Meox2-Cre^+/-^ mice show a typical DS-like phenotype^34,35^. Additionally, comparing C57Bl/6J WT and Meox2-Cre^+/-^ mice (Suppl. Fig. 2), we showed that Meox2-Cre^+/-^ mice do not have delay in the neurodevelopmental features that we have tested. All experiments were performed according to policies on the care and use of laboratory animals of the European Directive 2010/63/EU; animal experimental protocols have been validated by ethics committees (CIEPAL Azur) and the French Ministry of research under the agreement number APAFIS#9901-2017040616024590v6-PEA216/408 and APAFIS#15665-201805301624157v2-PEA499. Animals’ health was evaluated daily with a scale established a priori for signs of discomfort and pain (disclosed in the approved animal experimentation protocols). We have calculated a priori the number of mice to use per group with the software [G-Power](https://www.psychologie.hhu.de/arbeitsgruppen/allgemeine-psychologie-und-arbeitspsychologie/gpower), evaluating the variability observed in the experiments according to previous experience of the team in similar projects, preliminary experiments and reports in the literature, and taking into account the average mortality rate of *Scn1a*^+/-^ mice. Animals were randomized for allocation within each experimental group and the order of treatments and measurements were randomized with the online tool [Randomizer](http://www.randomizer.org/).

### Pharmacological treatments

KU55933 (Tocris, S1092) at the indicated concentration (7,5 mg/kg) diluted in the vehicle solution (DMSO) was delivered by intranasal administrations in a volume of 1 ml/kg, as in ^36^. Animals were treated every 72 hours from P15 to P21 (short treatment, two administrations) or until P30 (long treatment) (Fig. 1C, 3A, 4A). Bumetanide (Sigma, B3023) at the indicated concentration (0,2 mg/kg) diluted in the vehicle solution (PBS with 1% DMSO) was delivered with intraperitoneal daily injections in a volume of 0,02 ml/g from P15 for one or two weeks depending on the experiment (Fig. 3A & 4A). Similar Bumetanide treatments have been used in several in vivo studies of mouse models of brain diseases ^37–39^. For *ex-vivo* electrophysiology experiments, for which a single animal per day is used, we treated animals in sliding windows that allowed us to compare treated and controls animals from the same litter and to minimize the number of animals used for the study. We did not observe differences comparing recordings within the sliding window.

**Figure 1.**
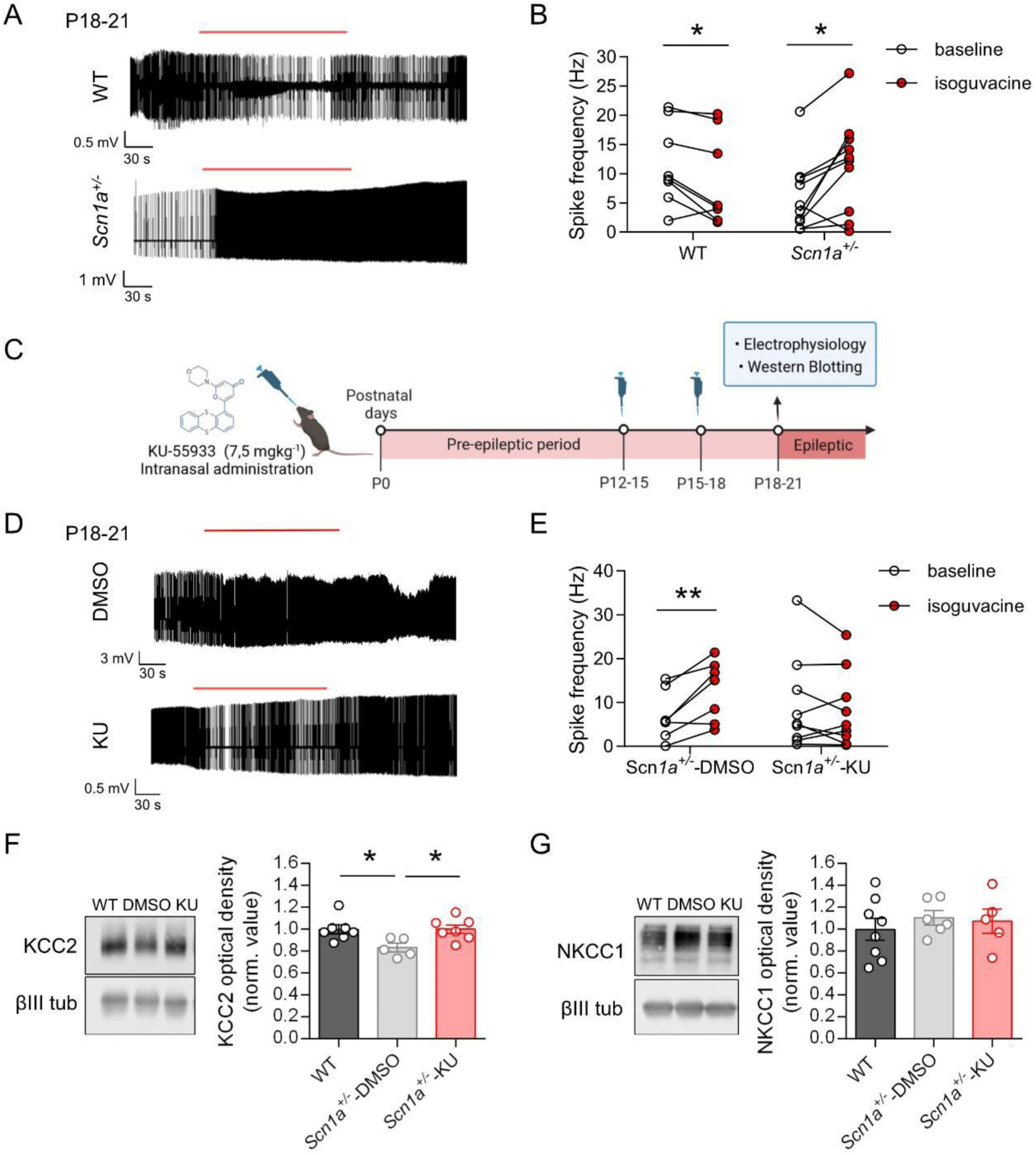
KU rescues the altered GABA switch in *Scn1a^+/-^* mice. **A.** Representative traces of CA3 pyramidal cells’ spontaneous spiking activity measured by current clamp (I=0) recordings in loose patch configuration. Cells were recorded for 3/5 minutes to obtain a stable control baseline, isoguvacine (10 µM) was bath applied for 3 minutes (bars above the traces) and washed out for at least 10 minutes. **B.** Analysis of mean spike frequency in control and upon isoguvacine application in WT or *Scn1a*^+/-^ CA3 pyramidal neurons revealed an excitatory effect of the GABA agonist in *Scn1a^+/-^* mice (paired t-test, p-value <0.05) whereas an inhibitory effect was observed in WT mice (paired t-test, p-value <0.05). **C.** Schematic representation of KU55933 (KU) treatment and experimental timeline. For comparing littermates in *ex-vivo* electrophysiological recordings, we used sliding windows: mice received two intranasal administrations (every 72 hours) of KU or vehicle (DMSO), the first in the window P12-P15 and the second in the window P15-18, and recordings were performed in the window P18-P21. For western blotting analysis, mice received two intranasal administrations (every 72 hours) at 15 and P18, and western blot was performed at P21. **D.** Representative traces of P18-21 CA3 pyramidal cells’ spiking activity and effect of isoguvacine (10 µM; bar above the traces) measured as in A. **E.** Analysis of mean spike frequency of baseline and upon isoguvacine application (bars above the traces) of *Scn1a*^+/-^-DMSO and *Scn1a*^+/-^-KU CA3 pyramidal neurons revealed an excitatory effect of GABA in vehicle-treated *Scn1a^+/-^* mice (paired t-test, p-value <0.01), whereas no significant modification was found in KU-treated animals between baseline and isoguvacine (paired t-test, p-value=0.4). **F.** Western blotting quantification of P21 hippocampal tissues showed a reduced expression of KCC2 chloride transporter in *Scn1a*^+/-^ mice, which is related to GABA switch delay. KU treatment rescued this alteration increasing KCC2 expression. **G.** There was no difference in NKCC2 levels in the three conditions. See Supplementary material for values, n and statistical tests.

### Preparation of brain slices and electrophysiological recordings

Procedures were similar to those reported in ^40,41^. Mice were deeply anesthetized with Isoflurane and decapitated. The brain was quickly removed and placed in ice-cold standard artificial cerebrospinal fluid (s-ACSF), which contained (in mM): 125 NaCl, 2.5 KCl, 2 CaCl_2_, 1 MgCl_2_, 1.25 NaH_2_PO_4_, 25 NaHCO_3_ and 25 glucose, saturated with 95 % O2 - 5 % CO_2_. Acute coronal slices (300 μm thick) were prepared with a vibratome (HM650V, MicroM, Germany) in ice-cold s-ACSF. Slices were then stored in a submerged chamber with s-ACSF at 34°C for at least one hour before the beginning of the recordings. Slices were visualized using infra-red differential interference contrast (DIC) microscopy (Nikon Eclipse FN1, Japan) equipped with a CCD camera (CoolSnap ES2, Photometrics, USA). Electrophysiological signals were recorded with a Multiclamp 700B amplifier (CV-7B headstage), a Digidata 1440A acquisition board and pClamp 10.3 software (Molecular Devices, USA). Recordings were performed at 34°C using an inline solution heater and a temperature controller (TC-344B, Warner Instruments, Hamden, CT, USA), while continuously perfusing the slices with s-ACSF bubbled with 95% O_2_/5% CO_2_ (pH 7.4).

Whole-cell patch-clamp recordings were performed in CA1 pyramidal neurons of the hippocampus using borosilicate glass pipettes containing (in mM): 120 K-gluconate, 15 KCl, 2 MgCl_2_, 0.2 EGTA, 10 HEPES, 20 P-Creatine, Na_2_-0.2 GTP, 2 Na_2_-ATP, 0.1 leupeptin, adjusted to pH 7.25 with KOH (3–5 MΩ resistance). For recordings of post synaptic currents (PSCs) QX314 1.5 mM was added to the internal solution in order to stabilize the membrane potential. Whole-cell access resistance (10-25 MΩ) was monitored and cells showing unstable access resistance (> 20 MΩ) were discarded. Spontaneous glutamatergic currents (sEPSCs) were measured at the reversal potential for GABA-A receptor mediated events (−60 mV) and spontaneous GABA-A receptor mediated currents (sIPSCs) were measured at the reversal potential of glutamatergic events (+10 mV). To be considered, sEPSCs/sIPSCs had to exceed a threshold of 4-times the standard deviation of the RMS noise evaluated on an eventless window of the trace. The E/I ratio was calculated by dividing sEPSCs and sIPSCs frequencies measured in the same neuron.

Juxtacellular-loose patch recordings were performed with the same pipettes used for whole cell experiments, but filled with s-ACSF, and action potentials were recorded in the I=0 mode. Cells were recorded for 3/5 minutes to obtain a stable baseline, isoguvacine (10 µM) was bath applied for 3 minutes and washed out for at least 10 minutes.

Data were not corrected for the liquid junction potential. Electrophysiological signals were filtered at 2 kHz and sampled at 10 kHz. Data were analyzed off-line (pClamp-10 software, Axon Instruments).

### Golgi staining

WT and *Scn1a*^+/-^ mice (males and females; P11 or P21) were deeply anesthetized with intra-peritoneal injections of Kétamine (100 mg/kg) et Xylasine (16 mg/kg) and intracardiacally perfused with saline (0,9% NaCl) to remove blood followed by 4% paraformaldehyde. The brain was removed and postfixed in 4% paraformaldehyde for 24 hours, then it was stained with Golgi-Cox solution as described in ^42,43^ with slight modifications. For the impregnation the brains were stored at room temperature in darkness for 14 days. After this period the Golgi-Cox solution was removed and replaced by 25% sucrose solution for at least 48 hours. Coronal sections of 100 μm thickness were obtained from the hippocampus using a vibratome (VT1000S, Leica, Germany) and collected in TBS. Sections were mounted onto gelatin treated or Superfrost Plus Micro slides using TBS and let them air dry for 24 hours. The sections were processed with ammonium hydroxide for 10 min, rinsed with distilled water, dehydrated with descending ethanol series, and mounted with a xylene-based medium. Spine density and morphology were evaluated on the secondary branches of apical dendrite of pyramidal neurons located in the CA1 subfield of the dorsal hippocampus. At least 20 neurons per mice were evaluated.

### Western Blotting

The general procedure was described in ^36^. Proteins were obtained starting from explanted brain tissues (hippocampus) homogenized in sample buffer (sodium dodecyl sulphate 1% (SDS), 62.5 mM Tris-HCl (pH 6.8), 290 mM sucrose) with Protease Inhibitor Cocktail and PhosSTOP (Roche). Protein content was assessed by Coomassie Protein Assay Kit (Thermo Fisher). Sample absorbance was read through a spectrophotometer (Eppendorf BioPhotometer plus) set to 550 nm. Protein content was assessed through a bovine serum albumin-based standard curve. Proteins were separated by SDS-PAGE electrophoresis, blotted, and incubated with primary antibody followed by HRP-conjugated secondary antibody (Jackson ImmunoResearch) and revealed by Pierce ECL Western Blotting Substrate (Thermo Fisher Scientific) through Fusion FX Vilber. βIII tubulin was used as normalizer. The following primary antibodies were used: rabbit anti-KCC2 (Millipore), mouse anti-NKCC1 clone T4 (Developmental Studies Hybridoma Bank) and mouse anti-βIII tubulin (Sigma-Aldrich).

### Seizure induction by hyperthermia

The animal was placed in a small incubator and the core temperature was continuously monitored using a rectal probe. Body temperature was increased 0.5 °C every 2 min as previously described ^44^ until either a behavioral seizure occurred, or a core body temperature of 42.5 °C was reached. The severity of seizures was quantified using a modified Racine’s scale ^45^: 0 - No behavioral response; 1-motionless starring (with orofacial automatism); 2 – Head nodding; 3 – Unilateral forelimb clonus; 4 - Bilateral forelimb clonus; 5 – Rearing and falling; 6 - Generalized tonic-clonic seizures. Thermal induction was performed in P21 Scn1a^+/-^ animals treated with KU or BUM or with their respective controls.

### Video analysis of spontaneous seizures

Starting from P15, the first day of treatment with KU or BUM or the respective controls, littermates were placed in standard cages with the mother and continuously monitored by CMOS video cameras (HDCVI, 2 Mpx, Sony exmor) equipped with infrared light and connected to a digital video recorder (DVR, HDVCI 1080P). Seizures frequency and clusters were measured with a noninvasive semi-automated detection of convulsive seizures from videos, based on an open-source package of two software programs that we developed for automated video acquisition (VASD) and semi-automated detection of seizures (SASDI) ^46^.

### Behavioral tests (performed at P21)

#### Social Interaction

The assay was performed as previously described ^47^. The three-chamber apparatus, slightly illuminated (30 lux), is a non-transparent Plexiglas box (30×60 cm) with two transparent partitions that make left, center, and right chambers (30×20 cm). In the first 10-min session, the test mouse was placed in the center of the three-chamber apparatus for habituation purposes. The mouse was allowed to freely explore each chamber containing one empty wire cage (11 cm high x10cm diameter). In the second 10-min session, the sociability phase, an age- and sex-matched mouse of the same strain that had never been exposed to the test mouse was placed in one of two wire cages in the side chambers. The location of the stranger mouse in the right or left chamber was alternated between trials. The empty wire cage in the opposite chamber served as the inanimate object. The test mouse was placed again in the center, and allowed to freely explore the chambers for 10 min. The time spent by the test mouse sniffing and interacting with each of the two wire cages was manually scored. The sociability index was calculated as the difference between the time spent exploring the stranger mouse and the empty cage divided by the total time of exploration of both cages.

#### Y maze spontaneous alterations

Spontaneous alternation was measured using a Y-shaped maze constructed with three symmetrical grey solid plastic arms at a 120-degree angle (26 cm length, 10 cm width, and 15 cm height) as previously described ^48^. Mice were individually placed at the end of one arm and were allowed to freely explore the three arms for 8 minutes. Arm entry was defined when all four limbs were within the arm. A triad was defined as a set of three arm entries, when each entry was in a different arm of the maze. Videos were analyzed with the ANY-maze video-tracking system (Stoelting Europe, Dublin, Ireland). The number of arm entries and the number of triads were recorded to calculate the alternation percentage (generated by dividing the number of triads by the number of possible alternations, total arms entries −2, expressed as percentage).

### Neurodevelopmental tests (performed from P7 to P14)

#### Righting reflex

The pup was placed on its back and the time required to recover to a normal position with the 4 paws touching the surface was measured manually. This reflex was evaluated at P7 and P10-11 using a cut-off of 30 seconds. The test reflects both motor coordination and vestibular function ^49^.

#### Forepaw grasping reflex

When a thin rod is placed against the palm of one paw with a light pressure, the paw flexed to grasp the rod. We assessed if this motor reflex was acquired for one or both forepaws on P7 and P10-11.

#### Negative geotaxis reflex

The pup was placed, head down, on a rough surface inclined at 15° ^49^. The time taken to turn 180° was measured manually on P10-P11 using a cut-off of 60s. As the righting reflex, geotaxis reflex assesses both motor coordination and vestibular function.

#### Gait

The pups were placed in the center of a 15 cm diameter circle and the time needed to move outside the circle task was measured on P10-P11 ^50^.

#### Olfactory motivation (nest-bedding/homing test)

WT and *Scn1a*^+/-^ mice were tested for olfactory motivation at P10-11 by placing them in the center of a square (7 cm × 7 cm) plastic box as in ^51^. One corner of the tray contained bedding from the home cage, while the opposite one contained new clean bedding. Mice were placed with their heads pointed toward an empty corner of the tray, forced to turn left or right to orientate toward the bedding corners. Mice were allowed up to one minute to reach a corner, before the trial was stopped. Trials in which mice failed to move, or to arrive at a bedding corner, were considered uncompleted trials, whereas those in which mice arrived at a bedding corner within one minute were considered completed. The latency to reach one of the bedding corners and the type of bedding corner reached were recorded for each of the 3 trials. In each trials the beddings were in different orientations relative to the mouse. The percentage of success was calculated as the percentage of trials in which the mice reached the home cage bedding on the total of 3 trials.

#### Eye opening

The number of open eyelids was checked at P13-P14.

### Data analysis and statistics

Data analysis was performed with pClamp 10 (Molecular Devices, USA), Excel (Microsoft Office, USA), ImageJ, Prism9 (GraphPad, USA) and Origin2021 (OriginLab, USA). Statistical tests were performed with Prism9 or Origin2021. The Fisher Exact test or the chi-square test were used for comparing categorical data. Other comparisons were performed with the Student’s t-test (two groups, paired or unpaired) and one-way ANOVA followed by Fisher’s LSD post-hoc test (three groups) (19) for normally distributed data, the Mann-Whitney test (two unpaired groups) and the Wilcoxon signed rank test (two paired groups) for non-normally distributed data. Normality was tested with the Kolmogorov-Smirnov (KS) test. The KS test was used also for comparing two empirical distributions. Differences were considered significant when P values where smaller than 0.05 (*P<0.05, **P<0.01, ***P<0.001, ****P<0.0001).

## Results

### GABA-switch is delayed in DS *Scn1a*^+/-^ mice

As DS model, we used F1 mice generated mating floxed homozygous *Scn1a* C57BL/6J females ^15,34^ and heterozygote Meox2-Cre C57BL/6J males (Jackson Laboratories, stock No: 003755) to obtain *Scn1a*^+/-^ mice. This strategy has been already used in previous studies and these *Scn1a*^+/-^ mice show a typical DS-like phenotype ^34,35^.

We evaluated the effect induced by the activation of GABA-A receptors on the spontaneous excitability of CA3 excitatory hippocampal pyramidal neurons in brain slices of P18-P21 *Scn1a*^+/-^ mice, an age at which the GABA-switch should be accomplished and before onset of spontaneous seizures in this model. Hippocampal CA3 neurons have been used in numerous studies that have evaluated the kinetics of GABA-switch ^29–33^. We performed current clamp (I=0) electrophysiological recordings of spontaneous spiking activity in loose patch configuration, quantifying the modifications of spiking activity induced by the application of the GABA-A receptor agonist isoguvacine (10 µM) ^52^. We found a preponderant depolarizing response leading to increased excitability in pyramidal neurons of *Scn1a*^+/-^ mice, whereas in WT littermates most of the cells showed a reduction of spike frequency, although the reduction did not reach statistical significance (Fig. 1A,B). These results confirm that the GABA-switch is delayed in DS mouse models, as it has been reported in a different DS *Scn1a* knock-out model measuring the reversal potential of GABA-evoked currents ^24^.

### KU55933 normalizes GABA-switch and decreases the excitatory/inhibitory ratio in *Scn1a^+/-^* mice

A delayed GABA-switch has been implicated in numerous neurodevelopmental diseases ^31–33,53^. We investigated its consequences on the overall phenotype and its importance as pathological mechanism in *Scn1a*^+/-^ DS mice. We used two different pharmacological strategies, targeting the two chloride transporters that are responsible of the intracellular chloride homeostasis in neurons: KCC2, involved in chloride efflux, and NKCC1, involved in chloride influx. We exploited KU55933 (KU) which, blocking ATM kinase activity, acts on the GABA-switch by increasing the expression of KCC2 ^36,54^, and bumetanide, a FDA-approved NKCC1 specific antagonist ^55–57^. The action of bumetanide has been reported in numerous studies, whereas KU is less characterized. Therefore, we investigated the effects of KU in *Scn1a^+/-^* mice performing *ex-vivo* electrophysiological experiments. We treated mice in the period preceding the onset of spontaneous seizures (equivalent to the febrile stage in patients) using intranasal delivery of KU (7,5 mg/kg) to efficiently cross the BBB and to target specifically and directly the central nervous system, as in ^36,58^. We performed the treatment (two administrations every 72 hours) with sliding windows starting from P12-P15 and the recordings at P18-21 (three days after the last dose of KU or DMSO); biochemical analysis was performed at P21 starting the treatment at P15 (Fig. 1C). The sliding windows allowed us to compare treated and control mice from the same litter and to minimize the number of animal used for electrophysiological recordings, which are performed with a single animal per day. The electrophysiological evaluation of the effect of the treatment was obtained measuring the effect induced by the application of the GABA-A receptor agonist isoguvacine on the spontaneous spiking activity of CA3 pyramidal cells in hippocampal slices, which showed a rescue of the delayed GABA-switch that we observed in control conditions in *Scn1a^+/-^* mice (Fig. 1D-E). Western blot experiments showed that the dysregulation of the GABA-switch in *Scn1a^+/-^* mice was correlated to decreased expression of KCC2, which was normalized by the treatment with KU (Fig. 1F). Conversely, we found no changes in NKCC1 expression, neither in control nor upon KU treatment (Fig. 1G).

Notably, besides the GABA-switch (which is a postsynaptic feature), KU acts also on the overall kinetics of maturation of synaptic transmission and thus on the overall features of the network ^36,54^. Thus, we evaluated the effect of KU on spontaneous GABAergic (sIPSCs) and glutamatergic (sEPSCs) post-synaptic currents in hippocampal CA1 pyramidal neurons of KU- or vehicle-treated *Scn1a^+/-^* mice, compared to untreated WT littermates (Fig. 2A). The properties of sIPSCs and sEPSCs depend on both synaptic properties and network excitability, allowing to evaluate the overall features of the network. We analyzed hippocampal CA1 pyramidal neurons because they have been investigated in numerous previous studies and are clearly involved in the pathophysiology of DS ^59,60^. In vehicle-treated *Scn1a^+/-^* mice, we observed a reduced frequency of sIPSCs (Fig. 2B), as already reported in numerous studies of *Scn1a* loss of function. We also observed an increased frequency of sEPSCs (Fig. 2C), possibly induced by the reduction of IPSCs, leading to an increase of the excitatory/inhibitory ratio (Fig. 2D). We observed a small increase of sIPSCs and sEPSCs amplitude analyzing cumulative distributions, which was not significant comparing mean values (Fig2E, F). The KU treatment decreased the frequency and the amplitude of sEPSCs (Fig. 2C and 2F), whereas we observed only minor effects on sIPSCs (Fig. 2B and 2E). Despite the small effect on sIPSCs, the KU treatment was sufficient to rescue the E/I ratio of the cells (Fig. 2D). Although this is an interesting result, it should be kept in mind that the E/I ratio is a general parameter, obtained from mean values, which does not reflect the actual dynamical relationship between excitation and inhibition in the neuronal network.

**Figure 2.**
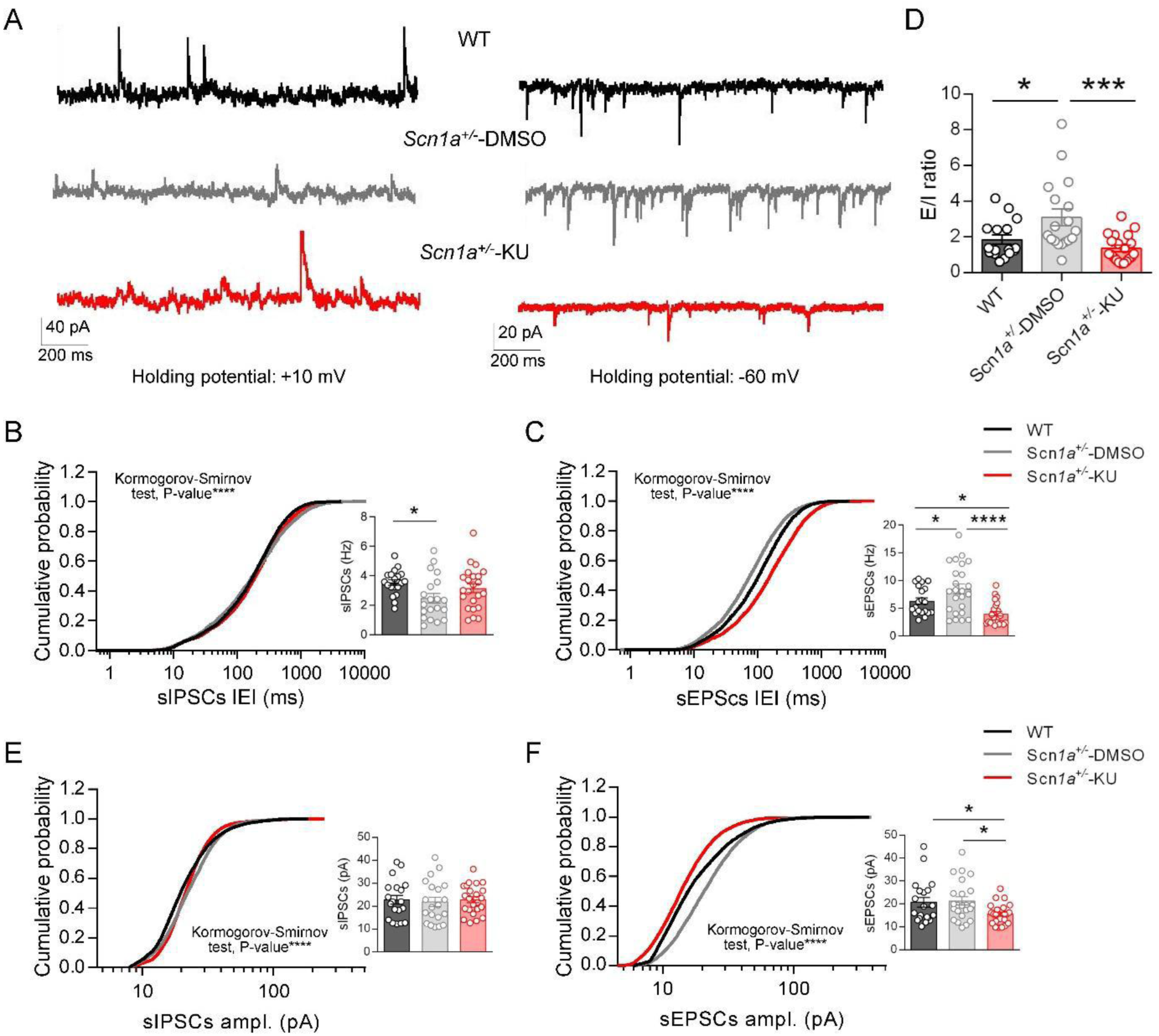
KU ex-vivo characterization in P18-21 *Scn1a^+/-^* mice: effect on spontaneous synaptic activity. **A.** Representative traces of spontaneous GABAergic post-synaptic currents (sIPSCs) and spontaneous glutamatergic post-synaptic currents (sEPSCs) recorded at +10 mV and −60 mV, respectively, in pyramidal neurons from hippocampal CA1 slices of vehicle-treated (DMSO) or KU-treated *Scn1a^+/-^* mice compared to WT littermates. **B.** The cumulative distribution showed an increased inter-event interval (IEI) of sIPSCs in vehicle-treated *Scn1a*^+/-^ mice compared to WT. KU treatment decreased sIPSCs IEI in *Scn1a*^+/-^ treated animals (KS test p-value <0.0001). Consistently, the comparison of mean sIPSCs frequency showed a decrease in vehicle-treated *Scn1a*^+/-^ mice compared to WT, which was partially rescued in KU-treated *Scn1a*^+/-^ mice. **C.** The cumulative distribution of sEPSCs IEIs displayed a decrease in vehicle-treated *Scn1a*^+/-^ mice and KU treatment rescued this defect, but increasing the IEI above WT level (KS test p-value <0.0001). Consistently, the comparison of mean sEPSCs frequency showed an increase in *Scn1a^+/-^* compared to WT and KU administration rescued this defect, but reducing the mean sEPSCs frequency below that of WT. **D.** The excitatory/inhibitory ratio (E/I) was obtained dividing mean sEPSCs and sIPSCs frequencies measured of the same neuron. **E.** The cumulative distribution of sIPSCs amplitude showed a slight difference between the distributions (KS test p-value <0.0001), which did not reach statistical significance comparing mean values. **F.** The cumulative distribution of sEPSCs amplitudes showed an increase in vehicle-treated *Scn1a^+/-^* mice and a decrease in KU-treated ones compared to WT (KS test p-value <0.0001). The decrease in vehicle-treated *Scn1a^+/-^* mice was not observed comparing mean values, whereas KU treatment decreased sEPSCs amplitude below WT level also considering mean values. See Supplementary material for values, n and statistical tests.

Overall, the KU treatment can rescue some of the dysfunctions that we observed, in particular the GABA-switch delay.

### Targeting GABA switch in *Scn1a^+/-^* mice has no effect on induced or spontaneous seizures

We performed *in vivo* experiments to evaluate the effect of KU and bumetanide on the epileptic phenotype of *Scn1a*^+/-^ DS mice, because the two treatments can rescue GABA-switch targeting different mechanisms. We treated animals starting from P15 (Fig. 3A) with intranasal administrations of KU (7,5 mg/kg every 72 hours) as in ^36^, or with bumetanide (2 mg/kg IP daily), method of administration that has been effectively used in numerous studies on different animal models of brain diseases ^37–39^. As for the *ex vivo* experiments, we began treatments at P15 to simulate clinical treatments, which do not start before the onset of the disease at the febrile stage. We initially evaluated the effect on hyperthermia-induced seizures at P21. Both treatments were ineffective in modifying the threshold temperature for seizure induction (Fig. 3B, D) or the severity of seizures, quantified using a modified Racine scale (Fig. 3C, E; see methods), which were similar to the values that we observed in control *Scn1a*^+/-^ mice treated with vehicle. To further characterize the contribution of the developmental delay linked to GABA-switch on the epileptic phenotype in *Scn1a*^+/-^ DS mice, we quantified in separated animal cohorts spontaneous convulsive seizures identified by semi-automatic analysis of high-resolution video recordings ^46^. This method allowed to avoid the surgical procedure for implanting electrodes for chronic recordings, which are difficult to implement at early ages, and permitted to chronically evaluate the animals for a long period starting from P15, the first day of the treatment. We performed video recordings for fifteen days (24/24h, 7/7d), from P15 to P30, on animals chronically treated with KU or bumetanide (Fig. 3A). Neither KU nor bumetanide reduced the frequency of seizures (Fig. 3F, I). We also analyzed the number of seizure clusters (we considered a cluster as two or more seizures with a delay of less than fifteen minutes) and found no changes in animals treated with KU or bumetanide compared to controls (Fig. 3G, J). Also, the two compounds had no effect on the survival rate of DS mice (Fig. 3H, K). Thus, neither KU nor bumetanide had effect on hyperthermia-induced or spontaneous seizures, showing that the long-lasting depolarizing effect of GABA does not play a crucial role on epileptogenesis or seizures generation in *Scn1a*^+/-^ DS mice.

**Figure 3.**
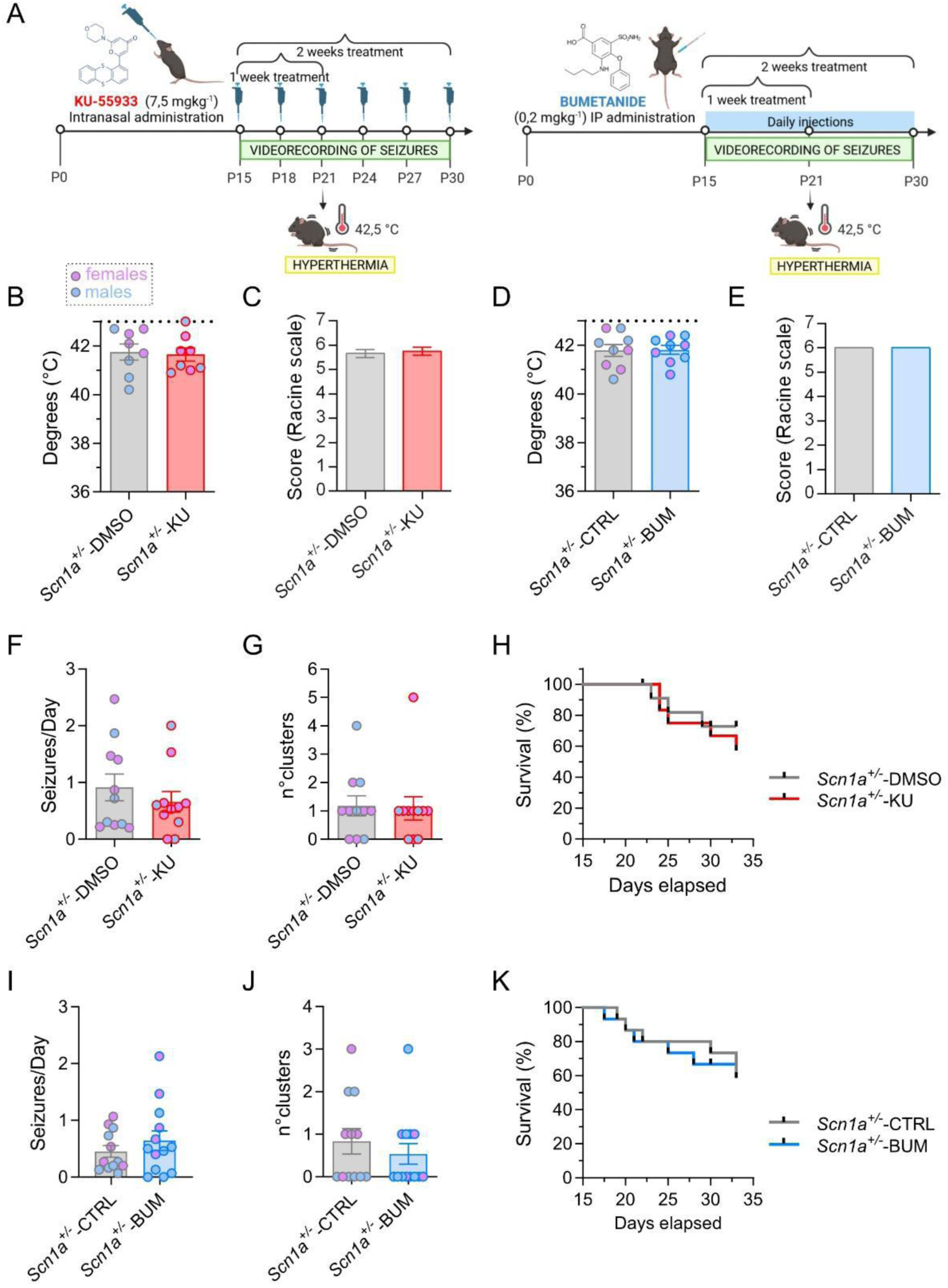
KU and bumetanide treatments do not affect the epileptic phenotype of *Scn1a*^+/-^ mice. **A.** Representative diagrams of KU55933 (KU) and Bumetanide (BUM) treatments and experimental timelines. Mice received KU 7.5 mg/kg intranasally every 72 hours or bumetanide 2 mg/kg IP daily starting from P15. The effect on hyperthermia induced seizures were assessed at P21 after a 1-week treatment. The effect on spontaneous seizures was assessed in separated animal cohorts treated chronically for two weeks. **B, C.** Hyperthermia-induced seizures did not show any differences upon KU treatment, neither in the threshold temperature nor in the seizures’ severity score. **D, E**. Similar to KU, BUM did not affect neither the threshold temperature nor the severity score. **F-G.** Video-analysis of spontaneous convulsive seizures showed no changes in KU-treated vs DMSO-treated mice in seizures’ frequency (F) or number of clusters of seizures (G). **H.** KU did not improve the mortality rate (p-value = 0.9755 Kaplan-Meier). **I-J.** Similar to KU, Bumetanide treatment did not improve seizures’ frequency (I) or clusters of seizures (J). **K.** BUM treatment did not modify the mortality rate (p-value = 0.2356; Kaplan-Meier). See Supplementary material for values, n and statistical tests.

### Rescuing the GABA-switch delay ameliorates sociability, hyperactivity, and cognitive deficits in juvenile *Scn1a^+/-^* mice

Since cognitive/behavioral dysfunctions are a major issue in DS, we investigated whether the pharmacological rescue by KU or bumetanide of the abnormal GABA-switch could ameliorate the cognitive and behavioral deficits of *Scn1a^+/-^*mice. We began the treatment at P15, and we performed behavioral tests at P21 (Fig. 4A). We analyzed juvenile animals before the establishment of a strong epileptic phenotype to discriminate precisely the neurodevelopmental component involved in behavioral features from secondary effects induced by seizures.

**Figure 4.**
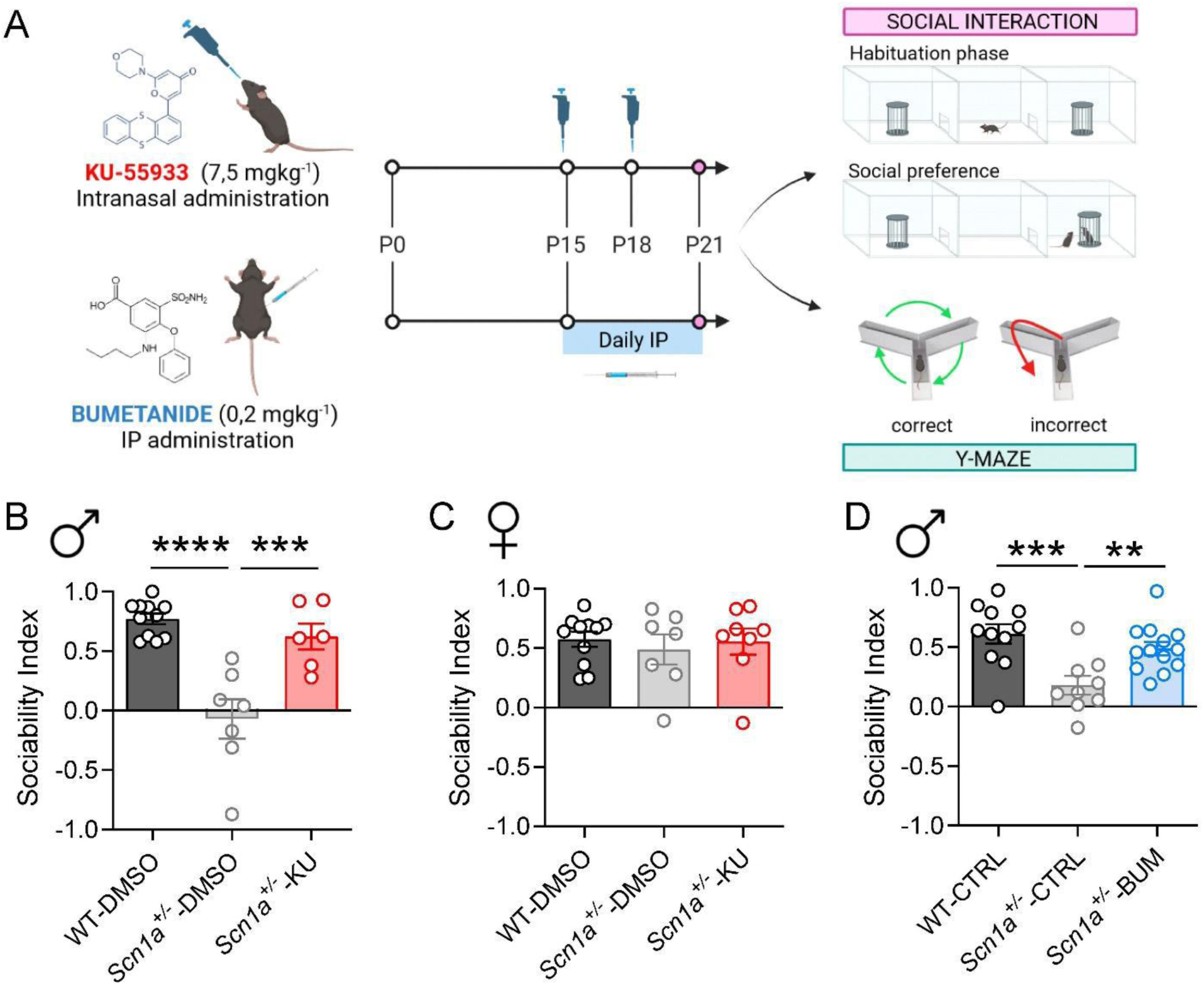
KU and bumetanide treatments ameliorate sociability of *Scn1a*^+/-^ mice. **A.** Diagram of the experimental timeline to test effects of KU55933 (KU) and bumetanide (BUM) on *Scn1a*^+/-^ autistic-like phenotype. Animals received KU treatment (two intranasal administrations, at P15 and P18, spaced of 72 hours) or BUM treatment (one-week daily IP injections from P15). Three-chamber sociability test and Y-maze were performed at P21. **B.** *Scn1a^+/^*^-^ male mice showed a defect in social preference compared to WT at P21, as indicated by a lower sociability index, and KU treatment rescued this alteration. **C.** The three-chamber test showed no defects in social interaction in *Scn1a^+/^*^-^ females. **D.** Like KU, BUM treatment rescued defective sociability in *Scn1a^+/^*^-^ male mice. See Supplementary material for values, n and statistical tests.

We initially focused on autism-like features, assessing deficits in social interactions, which have been already reported in adult *Scn1a*^+/-^ mice ^11,61^. In the three-chamber test, *Scn1a*^+/-^ male mice had profound defects in social interactions already at P21. Notably, targeting GABA-switch with KU or bumetanide rescued this behavioral defect. Both WT and *Scn1a*^+/-^ displayed no preference for the left or right chamber during the habituation period (Supplementary Fig. 1). However, when a stranger mouse was placed in one of the chambers, WT mice spent more time in close interaction with the stranger mouse than in the empty cage. Conversely, *Scn1a*^+/-^ mice had no preference for the stranger mouse. Remarkably, both KU and bumetanide treatments rescued the impairment in social interaction (Fig. 4B, D). As already reported for adult mice ^34^, juvenile *Scn1a*^+/-^ female mice did not show impairments in the three-chamber test (Fig. 4C).

Then, we tested cognitive features in juvenile *Scn1a*^+/-^ mice using the Y maze spontaneous alternation test, which can be used to assess spatial working memory. Spontaneous alternation is a behavior driven by the innate curiosity of rodents for exploring previously unvisited areas and can be evaluated by allowing mice to explore freely all three arms of the maze. Moreover, the number of entries in each arm and the total distance travelled can be used for evaluating locomotor (hyper-) activity. We tested P21 *Scn1a*^+/-^ mice treated with KU or bumetanide and compared them with WT littermates, considering males and females separately (Fig. 5). Both groups of control *Scn1a*^+/-^ male mice displayed a statistically significant decrease in the percentage of spontaneous alternations compared to WT males (Fig. 5A, D). In contrast, performances of *Scn1a*^+/-^ females were not different compared to WT female mice (Fig. 5G, J). Both male and female *Scn1a*^+/-^ DS mice showed a higher number of entries in the arms of the Y maze (Fig. 5B, E, H, K) and covered significantly longer distances during exploration (Fig. 5C, F, I, L). These data are in line with previous reports of hyperactive behavior in DS mouse models (52,51,70), and show that hyperactivity can be already observed in the pre-epileptic period. As shown in Fig. 5A, treatment with KU did not improve the dysfunctions of *Scn1a*^+/-^ males in spontaneous alternations. Although it did not reach statistical significance, KU treatment showed a loose trend towards the rescue of total distance travelled by males (5C) and a strong trend towards reduction of both the number of entries and the total distance travelled by females (Fig 5H, I). Notably, bumetanide rescued the reduced alternations, the number of entries and the total distance travelled of *Scn1a*^+/-^ males (Fig. 5D, E, F). In *Scn1a*^+/-^ female mice, bumetanide rescued the number of entries and partially improved the total distance travelled (Fig. 5K, L).

**Figure 5.**
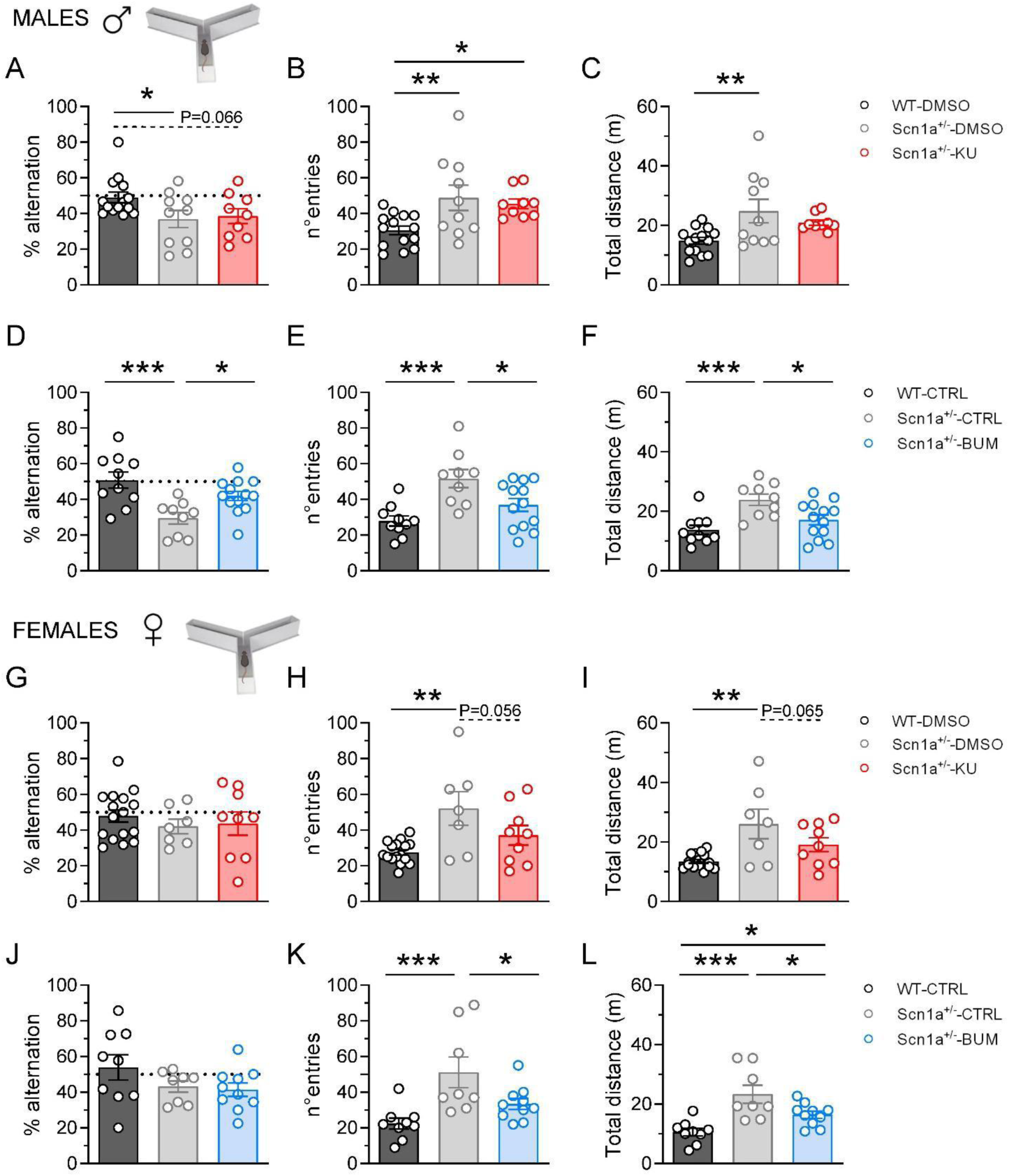
Bumetanide, but not KU, improves Y-maze performance and decreases hyperactivity in *Scn1a*^+/-^ mice. **A.** The Y-maze test showed a defect in spontaneous alternation in *Scn1a^+/^*^-^ male mice compared to WT at P21. KU treatment did not show a clear rescue of this defect. **B.** KU-treated and control DMSO-treated *Scn1a^+/^*^-^ male mice displayed an increased number of entries in the maze during the test. **C.** *Scn1a^+/^*^-^ male mice showed increased total distance compared to WT, and KU partially ameliorated this feature. **D.** The control group of *Scn1a^+/^*^-^ male mice used for the bumetanide (BUM) treatment showed, similarly to the DMSO-control ones used for testing KU, a defect in spontaneous alternation compared to WT, which was rescued by BUM. **E.** Control *Scn1a^+/^*^-^ male mice showed an increased number of entries which was rescued by BUM. **F.** Control *Scn1a^+/^*^-^ male mice showed an increased distance travelled, which was rescued by BUM. **G.** WT-DMSO, *Scn1a^+/-^*- DMSO and *Scn1a^+/^*^-^- KU female mice did not show differences in spontaneous alternation in the Y-maze test. **H.** Female *Scn1a^+/-^* mice, as male ones, showed an increased number of entries in the Y-maze, with a trend towards rescue upon KU treatment. **I.** Female *Scn1a^+/-^* mice, as male ones, showed increased total distance travelled during the Y-maze test, with a trend towards rescue upon KU treatment. **J.** As in the KU experimental cohort, Y-maze test showed no differences in Y-maze performance in WT, *Scn1a^+/-^*- CTRL and *Scn1a^+/^*^-^- CTRL female mice. **K.** BUM rescued the increased number of entries showed by *Scn1a^+/-^* mice during the Y-maze test. **L.** The increased total distance travelled by *Scn1a*^+/-^ mice during the Y-maze test was partially reduced by BUM, but it was still different from WT. See Supplementary material for values, n and statistical tests.

Therefore, targeting GABA-switch has an effect on autism-like features, spatial memory and hyperactive behavior in *Scn1a*^+/-^ mice.

### *Scn1a^+/-^* mice display early neurodevelopmental defects before seizures appearance

The timing of the GABA-switch is one of the features of GABA in neurodevelopment. In fact, GABA has trophic actions during early development before GABA-switch onset; its reduced release caused by Na_V_1.1 loss of-function may decrease depolarizing actions, which are important in these developmental stages (also for inducing the GABA-switch), and cause an overall delay in maturation ^31,32,63,64^. Thus, we investigated whether behavioral deficits in *Scn1a^+/-^* mice can be observed at early postnatal stages, when Na_V_1.1 is already expressed but before seizure onset. We quantified several developmental milestones, in particular those frequently used for characterizing dysfunctions in animal models of ASD ^51,65^. We analyzed motor ability and coordination at P7 and P10-11 and found no differences in righting reflex, grasping reflex, negative geotaxis, and gait (Fig. 6A-D). Olfactory detection and motivation to reach the nest and mother is one of the earliest behavioral correlates of social information processing, and a poor interest in reaching the home-cage bedding has been reported in mouse models of autism and socially anhedonic syndromes ^51,65,66^. We performed the nest bedding olfactory motivation test at P10-11. As shown in Fig. 6E, *Scn1a*^+/-^ mice displayed significant impairments in reaching the home cage bedding compared to WT, with no differences in the average time taken to reach the nest, consistent with no dysfunctions in sensory and motor abilities. Moreover, eye opening was delayed in *Scn1a*^+/-^ mice compared to WT at P13-14 (Fig. 6F), a feature that has been related to slowed and abnormal development of neuronal networks, also linked to anomalies in the development of the GABAergic system ^67,68^. To evaluate if Na_V_1.1 loss of function can reduce network activities also at these stages, we recorded GABAergic (sIPSCs) and glutamatergic (sEPSCs) post-synaptic currents in pyramidal cells of the CA1-area of hippocampal slices from P11-12 *Scn1a*^+/-^ mice (Fig.7A). We found a clear reduction in frequency (Fig.7C) and a mild reduction in amplitude of sIPSCs (Fig.7E) already at this early stage of development. Furthermore, we observed a mild reduction in the amplitude of sEPSCs, although their frequency was not modified (Fig. 2F).

**Figure 6.**
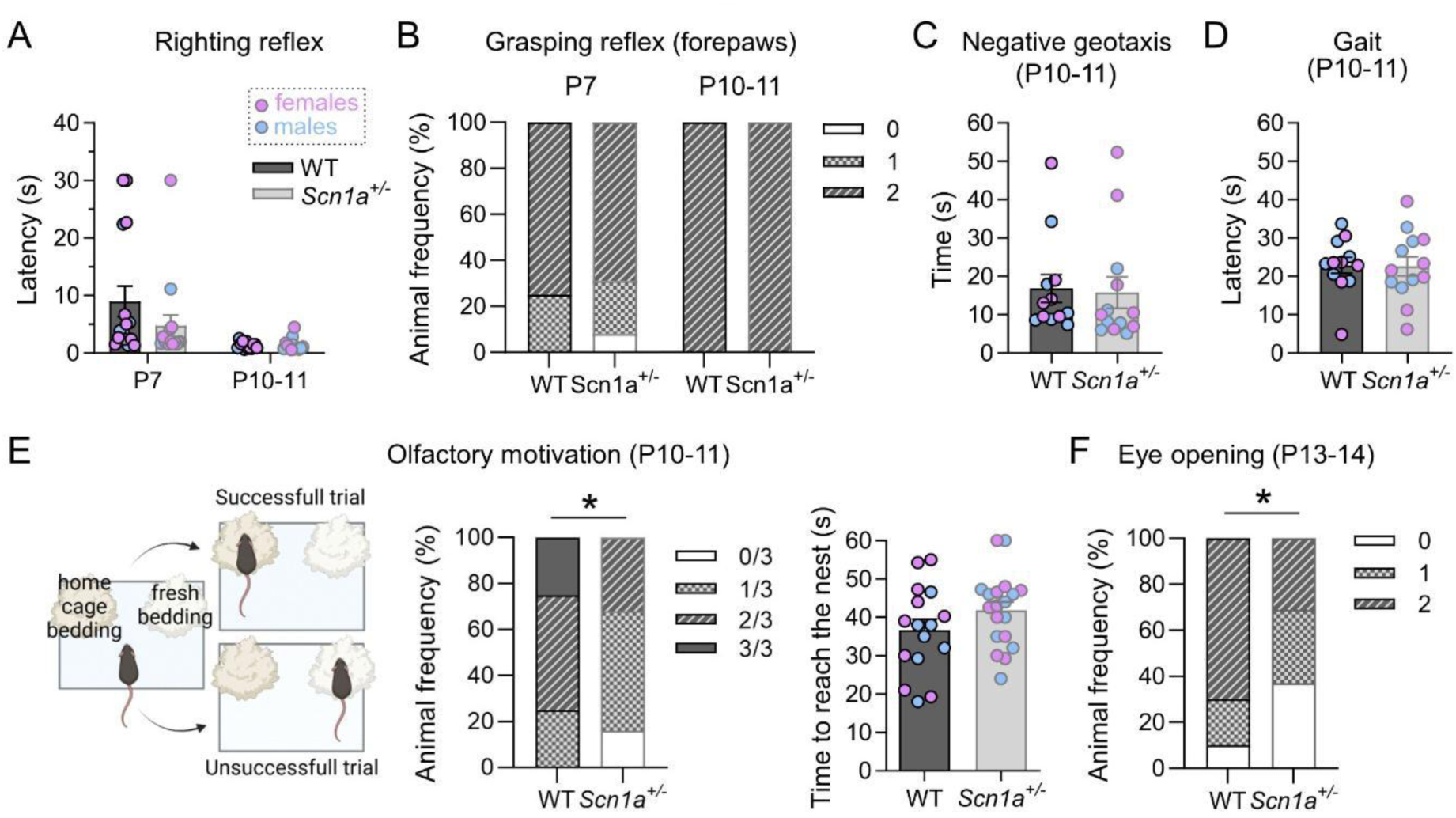
Early neurodevelopmental characterization of *Scn1a^+/-^* mice. **A.** Righting reflex analysis (cut-off 30 s) did not show any differences between WT and *Scn1a^+/-^* mice at P7 or P10-11. **B.** Grasping reflex test (forepaws) did not show any differences between WT and *Scn1a^+/-^*mice at P7 and P10-11. **C.** The negative geotaxis test at P10-11 gave similar results for *Scn1a^+/-^* and WT mice. **D.** The gait test showed no differences between *Scn1a^+/-^* and WT mice at P10-11. **E.** *Scn1a^+/-^* mice showed dysfunctions in the nest bedding test (olfactory motivation) compared with WT at P10-11. Three trials for each animal were performed and considered successful if the animal reached the nest bedding in less than 60 s. The time needed to reach the nest was similar for the two genotypes. **F.** Eye opening is delayed in *Scn1a^+/-^* compared with WT mice. See Supplementary material for values, n and statistical tests.

**Figure 7.**
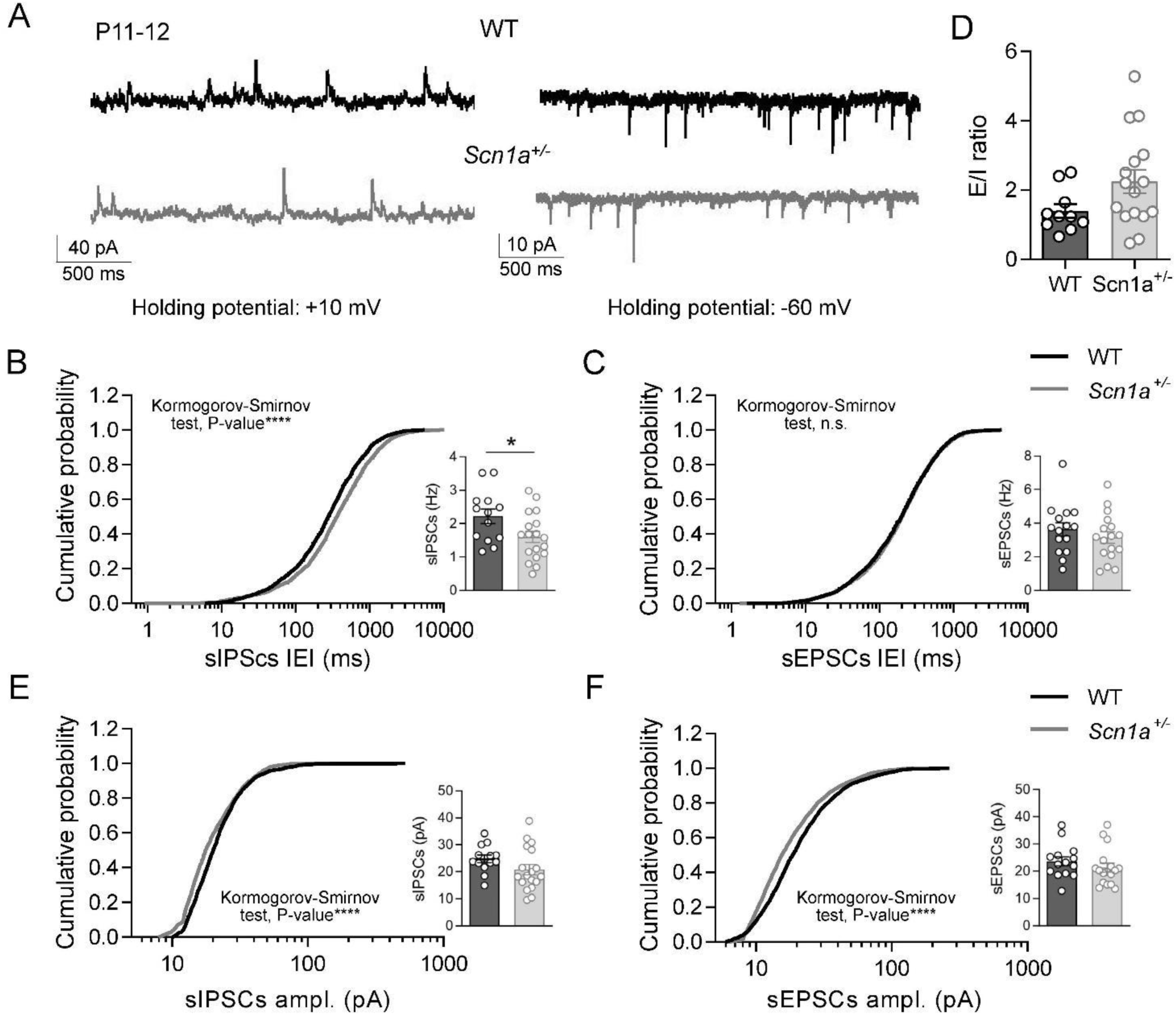
*Scn1a^+/-^* mice show defects in spontaneous synaptic activity before seizures appearance. **A.** Representative traces of spontaneous GABAergic post-synaptic currents (sIPSCs) and glutamatergic post-synaptic currents (sEPSCs) recorded in P10-12 hippocampal CA1 pyramidal neurons clamped at +10 mV and −60 mV, respectively. **B.** The cumulative distribution of sIPSCs inter event interval (IEI) revealed an increase in *Scn1a^+/-^* mice compared to WT, KS test p-value <0.0001. Consistently, the analysis of mean synaptic events showed a decreased frequency of sIPSCs in *Scn1a^+/-^* mice. **C.** Cumulative distribution of sEPSCs IEI revealed no changes in *Scn1a^+/-^* compared to WT mice, KS test p-value = 0.3141. Similarly, there were no changes in sEPSCs mean frequency in *Scn1a^+/-^* mice. **D.** The excitatory/inhibitory ratio (E/I) has been obtained dividing sEPSCs and sIPSCs frequencies measured in the same neuron. **E.** The cumulative distribution of sIPSCs amplitudes revealed a slight decrease in *Scn1a^+/-^* mice, KS test p-value <0.0001, which however did not reach statistical significance comparing mean values. **F.** Cumulative distribution of sEPSCs amplitude revealed a decrease in *Scn1a^+/-^* compared to WT mice, KS test p-value <0.0001, which did not reach statistical significance comparing mean values. See Supplementary material for values, n and statistical tests.

Features of dendritic spines are hallmarks of neurodevelopment and KCC2, which we found downregulated in *Scn1a*^+/-^ mice, is implicated in spine formation and maturation ^69,70^. We evaluated the features of dendritic spines of pyramidal neurons in the CA1 area of the hippocampus by using Golgi staining (Fig.8 A-B) and indeed observed a decreased spine head width at both P11 and P21 in *Scn1a*^+/-^ mice (Fig.8C), whereas spine density was not modified (Fig.8D). This is consistent with an impairment of dendritic spine maturation.

**Figure 8.**
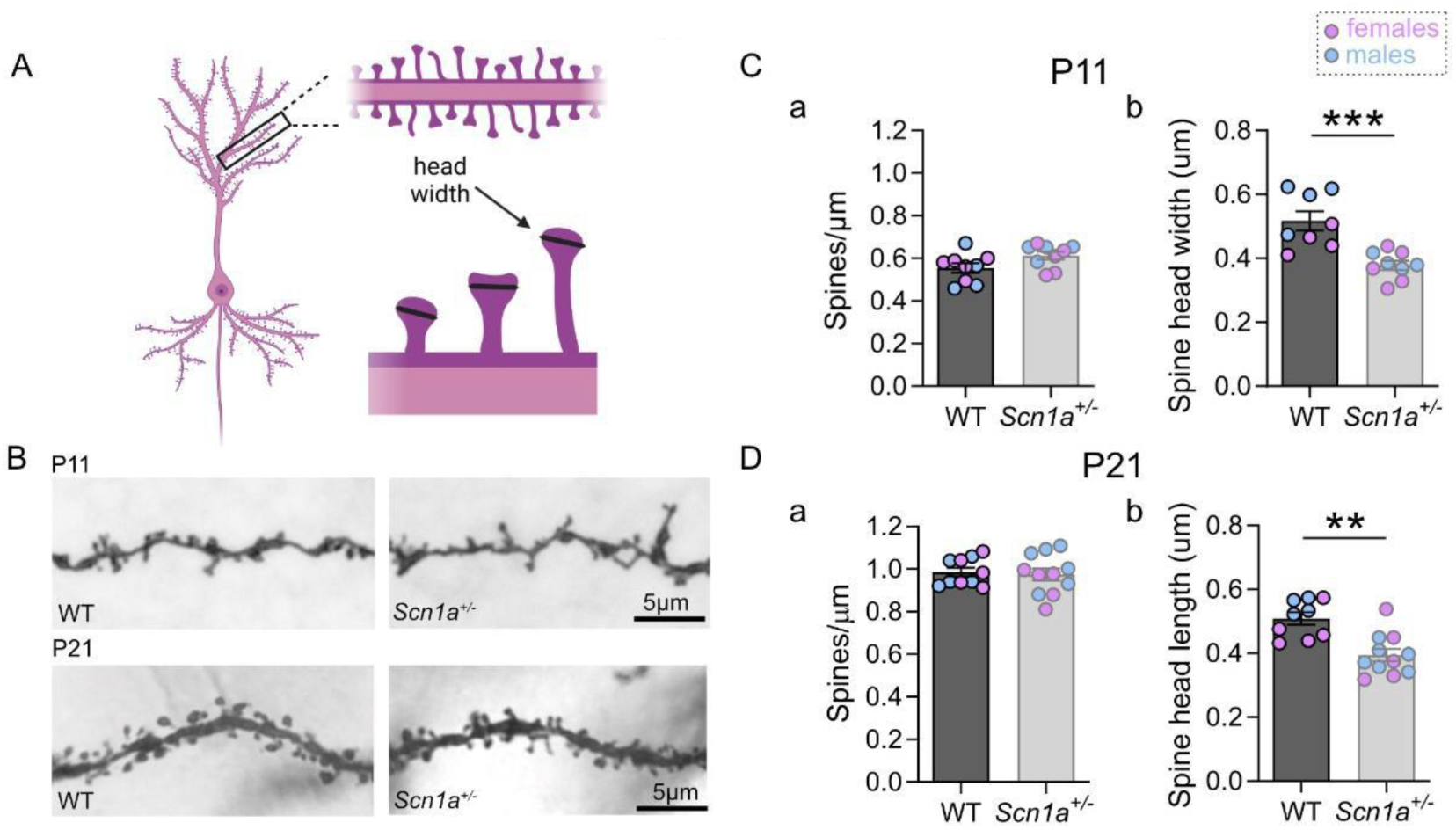
*Scn1a^+/-^* mice show defects in spine maturation. **A.** Representation of dendritic spines in CA1 pyramidal cell and illustration of how the morphometric parameter “head width” was quantified (black segments in the spine heads). **B.** Representative images of dendritic spines of pyramidal neurons in the CA1 area of the hippocampus of P11 (upper panels) and P21 (lower panels) in WT and *Scn1a*^+/-^ mice (Golgi staining). **C(a).** No change in spine density between WT and *Scn1a*^+/-^ P11 mice. **C(b).** Decreased spine head width in *Scn1a*^+/-^ mice at P11. WT and *Scn1a*^+/-^. **D(a).** No change in spine density in P21 *Scn1a*^+/-^ mice compared with WT. **D(b).** Decreased spine head width in *Scn1a*^+/-^ mice at P21. WT and *Scn1a*^+/-^. See Supplementary material for values, n and statistical tests.

Taken together these data show a neurodevelopmental component in *Scn1a*^+/-^ DS mice, with alterations appearing at early developmental stages at the beginning of Na_V_1.1 expression and before the onset of seizures, in which the decreased release of GABA and the consequences on its trophic actions may be involved.

## Discussion

The disease onset in DEEs is often considered as the onset of seizures, which is the initial most evident symptom of the pathology, but other dysfunctions, in particular cognitive/behavioral ones, may begin to develop earlier, and they are a persistent major burden in DS. Although numerous studies have reported in DS mouse models hypoexcitability of GABAergic neurons and reduced GABAergic synaptic transmission in the period preceding the onset of spontaneous seizures ^11–13,14(p1),15,18,71^, little is known about the link between these early dysfunctions of neuronal networks and of development of disease features, in particular of cognitive/behavioral defects. Notably, early pre-epileptic dysfunctions in the GABAergic system may lead to neurodevelopmental delay and contribute to the generation of neuropsychiatric symptoms ^30,64,72^. Interestingly, a delayed GABA-switch has been reported in models of DS ^24,25^, but its importance as a pathological mechanism in DS has not been investigated yet. A study showed that its pharmacological rescue with bumetanide was able to induce a slight delay of mortality in the *Scn1b^-/-^* mouse, which is however a model of sodium channel β1 subunit DEE ^24^, a different clinical entity with specific pathologic mechanisms and phenotype than DS ^1,26,27^.

Here we assessed the importance of the delay in the GABA-switch as a pathological mechanism in *Scn1a*^+/-^ mice, rescuing it by targeting either KCC2 or NKCC1 with KU55933 (KU) or bumetanide, respectively, and evaluating the effect on both epileptic and cognitive/behavioral features. For increasing the translational value of the results, we started the treatments at P15, an age that corresponds to the onset of the febrile stage in patients. Both drugs rescued social interaction deficits observed in P21 *Scn1a^+/-^* mice and ameliorated hyperactivity in the Y maze test; bumetanide also improved spatial working memory deficits. Although its blood-brain barrier permeability is low, similar bumetanide treatments have been effectively used in several *in vivo* studies of mouse models of neurodevelopmental diseases ^37–39,73^. In particular, our results are in agreement with previous studies showing beneficial effects of bumetanide treatment on hyperactivity and social behavior in other neuropsychiatric diseases: on hyperactivity in mice with dysfunctional K_V_7 voltage-gated K^+^ channels ^73^ and on both hyperactivity and social behavior in patients with tuberous sclerosis complex (TSC) ^74^. Thus, bumetanide, as well as other drugs targeting GABA-switch that may be approved in the future for clinical use (possibly including KU), could be beneficial also for DS patients. Conversely, neither KU nor bumetanide ameliorated the epileptic phenotype (hyperthermia-induced seizures, spontaneous seizures or mortality), although alterations of GABA-switch and KCC2 expression level/activity have been identified in epilepsy ^75^ and in our *ex vivo* experiments the KU treatment also decreased the frequency of spontaneous excitatory events in *Scn1a^+/–^* mice. This may be linked to the lack of effect of the treatment that we used on the reduced GABAergic synaptic transmission of *Scn1a*^+/-^ mice, which characterizes DS models and that is thought to be implicated in DS epileptogenesis. Taken together, our data suggest that GABA-switch delay is implicated in neuropsychiatric comorbidities of DS, in particular sociability defects (a prominent feature of ASD), but not in the epileptic phenotype.

Importantly, in addition to the delayed GABA-switch, we showed that DS mice display behavioral defects and delayed neurodevelopmental milestones at very early stages before the age of physiological GABA-switch onset, including olfactory discrimination deficits of nest bedding, which we observed at P10, and delayed eye opening. This is consistent with an early effect of the *Scn1a* mutation. In fact, we have shown that DS mice display a reduction of spontaneous GABAergic synaptic currents in CA1 pyramidal hippocampal neurons already at P11-12, an early developmental stage (well before seizure onset) in the first days after onset of Na_V_1.1 expression, which in mice is at around P8 ^16^. As already proposed in numerous previous studies ^11,12,14,76^, the reduced frequency of sIPSCs can directly depend on the Na_V_1.1 loss-of-function induced by DS mutations, which leads to a reduction of sodium current and excitability of GABAergic interneurons. However, this initial dysfunction of the GABAergic system could induce also a more general maturation delay. In fact, GABA exerts a trophic action at early stages of neurodevelopment promoting cell survival differentiation and migration, expression of brain-derived neurotrophic factor (BDNF), and spine maturation ^69,77,78^. Notably, GABA itself can promote GABA-switch ^79^ and reduced GABAergic transmission in *Scn1a*^+/-^ mice may interfere with it. Delayed GABA-switch might also be interpreted as a homeostatic response triggered by a reduction of the depolarizing action of GABA in DS models (because of hypoexcitability of GABAergic neurons) that leads to an overcompensation. However, delayed GABA-switch has been observed in numerous models of neuropsychiatric diseases and should not be specifically linked to the reduction in GABAergic transmission observed in DS models. Overall, because of its trophic role in neurodevelopment, the decreased GABA release in *Scn1a^+/-^* mice might be implicated in the behavioral defects, reduced amplitude of sEPSCs, and modifications of spine morphology that we have observed already at early developmental stages (P11-12). Interestingly, beside reducing GABAergic transmission, Na_V_1.1 loss-of-function can modify developmental activity patterns of neuronal networks possibly interfering with neuronal maturation ^63,80^. Overall, in the disease trajectory of DS, the GABAergic interneuron hypo-activity induced by Na_V_1.1 loss-of-function, which also reduces GABA release, should be the initial main event upstream to a series of further pathological dysfunctions, including the GABA-switch delay.

Our functional data correlates with the results of our western blot experiments, which showed decreased expression of KCC2 in the hippocampus of *Scn1a^+/-^* mice. This is at odds with the results reported for another Scn1a^+/-^ model that showed no modification of KCC2 expression, although a delay in GABA-switch was observed ^24^. However, that study used total brain membranes for western blot and a mixed 129×C57BL/6J genetic background that gives milder cellular dysfunctions and phenotype compared to the pure C57BL/6J mice that we used ^12,81^, which may explain the different results.

Features of dendritic spines are a hallmark of neurodevelopment and KCC2, independently of its chloride transport function, plays an important role in the development and morphology of dendritic spines through interaction with post-synaptic proteins and signaling pathways, leading to a remodeling of the spine’s actin cytoskeleton ^32,69,70^. This function could be implicated in the reduced spine head size without reduction of spine density that we observed in *Scn1a*^+/-^ mice. Consistently, KCC2 knock-out mice display spines with immature morphology (no obvious spine heads), whereas the density of the dendritic protrusions is not affected ^82^. Thin filopodia-like spine heads are considered less mature, indicating early stages of synapse formation or a lack of synaptic stabilization. Notably, altered spine morphology, including reduced head width, has been associated with pathological conditions, in particular neurodevelopmental diseases ^83^. Notably, also other dysfunctions that we have observed in *Scn1a*^+/-^ mice (i.e. E/I imbalance, defective GABAergic transmission, delay in neurodevelopmental milestones such as eye opening and GABA-switch) are key features observed in animal models of neurodevelopmental disorders, for example Down syndrome, Rett syndrome, Fragile X syndrome and the VPA model of autism spectrum disorder ^31,36,84–86^.

Our data showing the effects on behavioral dysfunctions, but not on seizures, of treatments targeting the delayed GABA-switch, as well as those showing early neurodevelopmental defects before seizure onset, demonstrate that DS mice display a neurodevelopmental delay that is specifically linked to the cognitive/behavioral phenotype, but not the epileptic phenotype. Thus, we disclose a neurodevelopmental component in DS which selectively impacts the behavioral phenotype, in particular autistic-like traits. We provide here new experimental evidence to the hypothesis that cognitive/behavioral defects in DS can develop before the onset of spontaneous seizures (as observed for some motor features and hyperactivity in the Scn1a^A1783V/+^ knock-in mouse model ^87^), and that they can be uncoupled from seizures and have specific pathological mechanisms, at least at the early stages of the disease. This can be relevant in a translational perspective. In fact, DS treatments are currently started after seizure onset and are mainly targeted to alleviation of seizures, whereas there are still no effective treatments against the cognitive/behavioral comorbidities, which are among the most severe phenotypic features, especially at later stages of the disease. Our results suggest novel therapeutic options for DS that can target cognitive/behavioral defects separately from seizures with specific pharmacological interventions that should be started as early as possible. In particular, targeting the GABA-switch may be an innovative pharmacological strategy to ameliorate DS behavioral defects.

Overall, our results are consistent with partially specific pathological mechanisms for seizures and some cognitive/behavioral defects in DS. This is a novel major finding for DS and DEEs. The early reduction of activity of GABAergic interneurons and the consequent reduction of GABA release in DS could determine a cascade of events resulting in abnormal neurodevelopmental trajectories, including delayed GABA-switch, because of the role of neuronal activity and GABA in neurodevelopment. These dysfunctions may be specifically targeted in therapy.

## Supporting information

Main Manuscript

## Acknowledgments

We thank Emilie Bonnet (Institute of Molecular and Cellular Pharmacology, CNRS UMR 7275 and University Cote d’Azur) for her skillful technical support.

This work was funded by UCAJEDI (https://univ-cotedazur.fr/ucajedi-lidex-duniversite-cote-dazur, ANR-15-IDEX-01, France), Laboratory of Excellence “Ion Channel Science and Therapeutics” - LabEx ICST (https://www.labex-icst.fr/en, ANR-11-LABX-0015-01, France) and EJPRD *SCN1A*-UP! (ANR-20-RAR4-0001, France) to MM. LP received a postdoc fellowship from the Fondazione Umberto Veronesi, Italy. FA was supported by Italian Ministry of University and Research, PRIN: Progetti di ricerca di rilevanza nazionale, 2022 (https://prin.mur.gov.it/Pages/Index/119) Prot.20229XWM5L, and PRIN: Progetti di ricerca di rilevanza nazionale, 2017 Prot. 20175C22WM.

## Author contributions

LP: designing research studies, conducting experiments, acquiring data, analyzing data, interpreting data, writing the manuscript.

FC: conducting experiments, acquiring data, analyzing data.

ER: conducting experiments, acquiring data, analyzing data.

GF: conducting experiments, acquiring data, analyzing data.

EM: analyzing data, interpreting data.

IL: conducting experiments, acquiring data, analyzing data, interpreting data.

FA designing research studies, interpreting data.

MM: designing research studies, analyzing data, interpreting data, writing the manuscript, supervising the project, acquiring funding.

