## Supplementary material for "Neurodevelopmental defects in Dravet syndrome *Scn1a*^+/-^ mice: targeting GABA-switch rescues behavioral dysfunctions but not seizures and mortality": Main Manuscript

### Supplementary figures

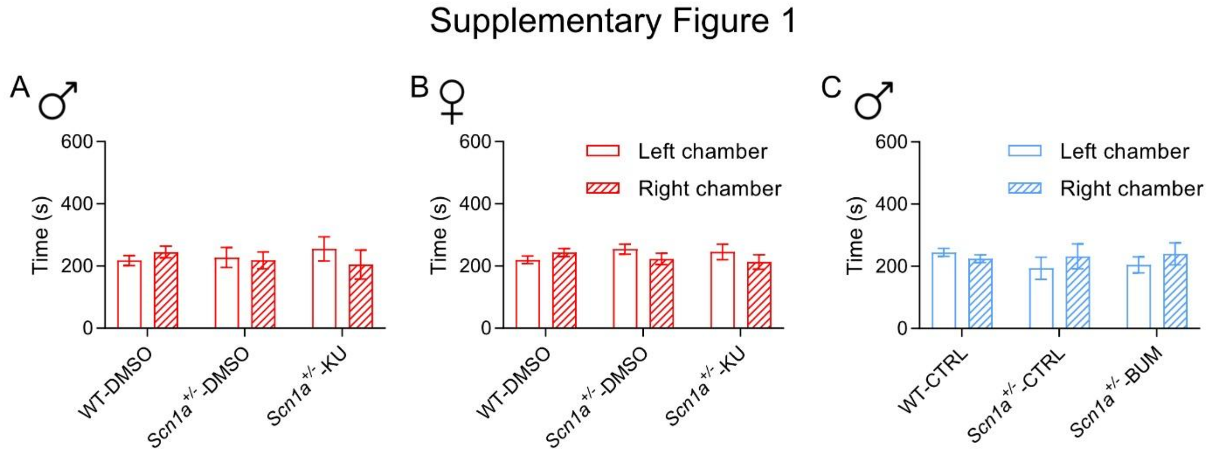

**Supplementary Figure 1. No preference for the left or right chamber during the habituation period of the three-chambers social interaction test.**

**A.** WT and Scn1a$^{+/-}$ KU-treated and non-treated male mice displayed no preference for the left or right chamber during the habituation period of the three-chambers social interaction test (WT-DMSO left: mean ± SEM = 217.545 ± 16.362, right: mean ± SEM = 244.764 ± 18.759, N = 11; Scn1a$^{+/-}$

- DMSO left: mean ± SEM = 227.429 ± 32.400, right: mean ± SEM = 218.514 ± 27.109, N = 7; Scn1a$^{+/-}$ - KU left: mean ± SEM = 220.120 ± 20.515, right: mean ± SEM = 245.600 ± 27.726, N =6; Multiple Wilcoxon test, p-value: WT-DMSO = 0.519531, Scn1a$^{+/-}$-DMSO = 0.687500, Scn1a$^{+/-}$-KU = >0.999999). **B.** As for males, WT and Scn1a$^{+/-}$ KU-treated and non-treated female mice showed no differences in the time spent in the left and rights chambers during the habituation phase (WT-DMSO left: mean ± SEM = 219.864 ± 12.359, right: mean ± SEM = 243.355 ± 13.146, N = 11; Scn1a$^{+/-}$ - DMSO left: mean ± SEM = 254.414 ± 16.112, right: mean ± SEM = 223.114 ± 18.130, N = 7; Scn1a$^{+/-}$ - KU left: mean ± SEM = 245.125 ± 25.008, right: mean ± SEM = 212.663 ± 23.436, N = 8; Multiple Wilcoxon test, p-value: WT-DMSO = 0.365234, Scn1a$^{+/-}$-DMSO = 0.296875, Scn1a$^{+/-}$-KU = 0.460938). **C.** WT and Scn1a$^{+/-}$ BUM-treated and non-treated male mice displayed no preference for the left or right chamber during the habituation period of the three-chambers social interaction test (WT-CTRL left: mean ± SEM = 242.550 ± 13.98, right: mean ± SEM = 230.910 ± 11.747, N = 10; Scn1a$^{+/-}$- CTRL left: mean ± SEM = 184.767 ± 23.478, right: mean ± SEM = 300.667 ± 23.102, N = 6; Scn1a$^{+/-}$- BUM left: mean ± SEM = 221.221 ± 27.804, right: mean ± SEM = 225.644 ± 46.408, N = 9; Multiple Wilcoxon test, p-value: WT-CTRL = 0.577148, Scn1a$^{+/-}$-CTRL = 0.546875, Scn1a$^{+/-}$-BUM = 0.839355).

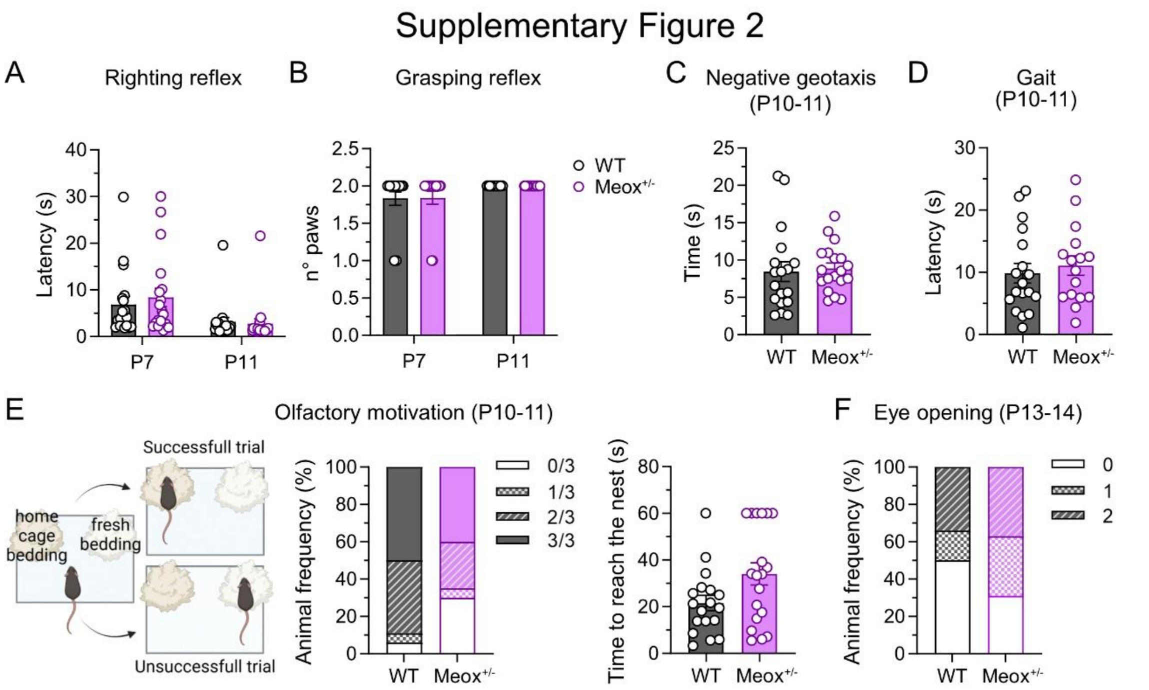

**Supplementary Figure 2. Meox$^{+/-}$ mice do not show delay in neurodevelopmental features**

**A.** Righting reflex analysis (cut-off 30 s) did not show any differences between WT and Meox$^{+/-}$ mice at P7 (WT: mean ± SEM = 7.180 ± 1.776, N = 17; Meox$^{+/-}$: mean ± SEM = 8.300 ± 1.986, N = 19) or P10-11 (WT: mean ± SEM = 3.114 ± 0.9851, N = 18; Meox$^{+/-}$: mean ± SEM = 2.743 ± 1.058, N = 19); Mann-Whitney test, p-value: P7 = 0.7844, P10-11 = 0.0714. **B.** Grasping reflex test (forepaws) did not show any differences between WT and Meox$^{+/-}$ mice at P7 (WT, N=18; Meox$^{+/-}$, N = 19) or P10-11 (WT, N = 18; Meox$^{+/-}$, N = 19); Fisher's exact test, p-value: P7 = >0.9999, P10-11 = >0.9999. **C.** The negative geotaxis test at P10-11 gave similar results for Meox$^{+/-}$ and WT mice (WT: mean ± SEM = 8.467 ± 1.315, N=18; Meox$^{+/-}$: mean ± SEM = 8.877 ± 0.7099, N = 19; Mann-Whitney test, p-value = 0.3090). **D.** The gait test showed no differences between Meox$^{+/-}$ and WT mice at P10-11 (WT: mean ± SEM = 9.858 ± 1.567, N = 18; Meox$^{+/-}$: mean ± SEM = 11.08 ± 1.575, N = 16; Mann-Whitney test, p-value = 0.5281). **E.** The nest bedding test (olfactory motivation) did not show differences between Meox$^{+/-}$ and WT at P10-11. Three trials for each animal were performed and considered successful if the animal reached the nest bedding in less than 60 s. The number of animals succeeding in 0, 1, 2 or 3 trials were compared between WT (N = 18) and Meox$^{+/-}$ (N = 19) mice; Fisher's exact test, p-value = 0.2604. The time needed to reach the nest was similar for the two genotypes; (WT: mean ± SEM = 21.53 ± 3.293; Meox$^{+/-}$: mean ± SEM = 34.05 ± 4.789); Mann-Whitney test, p-value = 0.0743. **F.** Eye opening is comparable between Meox$^{+/-}$ (N = 19) and WT (N = 18) mice (number of animals with 0, 1 or 2 open eyes at P13-14); Fisher's exact test, p-value = 0.4992.

### Statistical tables: values, n and statistical tests for main figures

##### Figure 1

| **Panel 1B** | WT baseline | WT isoguvacine | Paired t test | *Scn1a*$^{+/-}$ baseline | *Scn1a*$^{+/-}$ isoguvacine | Paired t test |
|---|---|---|---|---|---|---|
| Spike frequency (Hz) mean ± SEM | 11.58 ± 2.447 | 8.678 ± 2.740 | p=0.0230 n = 8 cells, N = 4 mice (2 males, 2 females) | 6.09 ± 1.94 | 11.52 ± 2.58 | p=**0.0129** n = 10 cells, N = 3 mice (2 males, 1 female) |

| **Panel 1E** | *Scn1a*$^{+/-}$-DMSO baseline | *Scn1a*$^{+/-}$-DMSO isoguvacine | Paired t test | *Scn1a*$^{+/-}$ KU baseline | *Scn1a*$^{+/-}$-KU isoguvacine | Paired t test |
|---|---|---|---|---|---|---|
| Spike frequency (Hz) mean ± SEM | 6.97 ± 2.13 | 12.69 ± 2.62 | p=0.009 n=7 cells, N = 3 mice (2 males, 1 female) | 9.51 ± 3.56 | 8.32 ± 2.89 | p=0.405 n=9 cells, N = 4 mice (3 males, 1 female) |

| **Panel 1F** | WT | Scn1a$^{+/-}$-DMSO | Scn1a$^{+/-}$-KU | One-Way ANOVA, Fisher's LSD post-hoc test |
|---|---|---|---|---|
| WB KCC2 normalized optical density mean ± SEM | 1.000 ± 0.041 N = 7 mice (4 males, 3 females) | 0.833 ± 0.038 N = 5 mice (3 males, 2 females) | 1.001 ± 0.037, N = 7 mice (4 males, 3 females) | p-value: WT vs Scn1a$^{+/-}$-DMSO = **0.0111**, WT vs. Scn1a$^{+/-}$-KU = 0.9789, Scn1a$^{+/-}$-DMSO vs. Scn1a$^{+/-}$-KU = **0.0106** |

| **Panel 1E** | WT | Scn1a$^{+/-}$-DMSO | Scn1a$^{+/-}$-KU | One-Way ANOVA, Fisher's LSD post-hoc test |
|---|---|---|---|---|
| WB NKCC1 normalized optical density mean ± SEM | 1.000 ± 0.098, N = 8 mice (5 males, 3 females) | 1.104 ± 0.065 N = 6 mice (3 males, 3 females) | 1.073 ± 0.117 N = 5 mice (2 males, 3 females) | p-value: WT vs Scn1a$^{+/-}$-DMSO = 0.4321, WT vs. Scn1a$^{+/-}$-KU = 0.5982, Scn1a$^{+/-}$-DMSO vs. Scn1a$^{+/-}$-KU = 0.8344 |

#### Figure 2

| **Panel 2B** | WT | $Scn1a^{+/-}$-DMSO | $Scn1a^{+/-}$-KU | One-Way ANOVA, Fisher's LSD post-hoc test |
|---|---|---|---|---|
| sIPSC frequency (Hz) mean ± SEM | 3.54 ± 0.19 n = 19 cells N=6 mice (3 males, 3 females) | 2.48 ± 0.33 n = 20 cells, N = 8 mice (3 males, 5 females) | 3.15 ± 0.29 n = 23 cells, N = 6 mice (3 males, 3 females) | p-value: WT vs $Scn1a^{+/-}$-DMSO = **0.0130**, WT vs. $Scn1a^{+/-}$-KU = 0.3270, $Scn1a^{+/-}$-DMSO vs. $Scn1a^{+/-}$-KU = 0.0978 (3 males, 3 females) |
| **Panel 2C** | WT | $Scn1a^{+/-}$-DMSO | $Scn1a^{+/-}$-KU | One-Way ANOVA, Fisher's LSD post-hoc test |
| sEPSC frequency (Hz) mean ± SEM | 6.30 ± 0.56 n = 20 cells, N = 6 mice (3 males, 3 females) | 8.45 ± 0.89 n = 24 cells, N = 8 mice (3 males, 5 females) | 3.99 ± 0.41 n = 24 cells, N =6 mice (3 males, 3 females) | p-value: WT vs $Scn1a^{+/-}$-DMSO = **0.0286**, WT vs. $Scn1a^{+/-}$-KU = **0.0190**, $Scn1a^{+/-}$-DMSO vs. $Scn1a^{+/-}$-KU **<0.0001** |
| **Panel 2D** | WT | $Scn1a^{+/-}$-DMSO | $Scn1a^{+/-}$-KU | One-Way ANOVA, Fisher's LSD post-hoc test |
| E/I ratio mean ± SEM | 1.85 ± 0.26, n = 16 cells, N = 6 mice (3 males, 3 females) | 3.09 ± 0.47, n = 18 cells, N = 8 mice (3 males, 5 females) | 1.36 ± 0.17, n = 20 cells, N = 6 mice (3 males, 3 females) | p-value: WT vs $Scn1a^{+/-}$-DMSO = **0.0108**, WT vs. $Scn1a^{+/-}$-KU = 0.2951, $Scn1a^{+/-}$-DMSO vs. $Scn1a^{+/-}$-KU = **0.0003** |
| **Panel 2E** | WT | $Scn1a^{+/-}$-DMSO | $Scn1a^{+/-}$-KU | One-Way ANOVA, Fisher's LSD post-hoc test |
| sIPSC amplitude (pA) mean ± SEM | 22.85 ± 1.96, n = 19 cells, N = 6 mice (3 males, 3 females) | 21.87 ± 2.06, n = 20 cells, N = 8 mice (3 males, 5 females) | 22.86 ± 1.32 n = 23 cells, N = 6 mice (3 males, 3 females) | p-value: WT vs $Scn1a^{+/-}$-DMSO = 0.7050, WT vs. $Scn1a^{+/-}$-KU = 0.9962, $Scn1a^{+/-}$-DMSO vs. $Scn1a^{+/-}$-KU = 0.6881. |
| **Panel 2F** | WT | $Scn1a^{+/-}$-DMSO | $Scn1a^{+/-}$-KU | One-Way ANOVA, Fisher's LSD post-hoc test |
| sEPSC amplitude (pA) mean ± SEM | 20.79 ± 2.10 n = 23 cells, N = 6 mice (3 males, 3 females) | 21.37 ± 1.80 n = 24 cells, N = 8 mice (3 males, 5 females) | 15.57 ± 0.90 n = 20 cells, N = 6 mice (3 males, 3 females) | p-value: WT vs $Scn1a^{+/-}$-DMSO = 0.8057, WT vs. $Scn1a^{+/-}$-KU = **0.0281**, $Scn1a^{+/-}$-DMSO vs. $Scn1a^{+/-}$-KU = **0.0119**. |

#### Figure 3

| **Panel 3B** | *Scn1a*$^{+/-}$DMSO | *Scn1a*$^{+/-}$ KU | Mann-Whitney test |
|---|---|---|---|
| Threshold temperature (°C) mean ± SEM | 41.75 ± 0.33°C N=8 mice (4 males, 4 females) | 41.65 ± 0.26°C N=8 mice (4 males, 4 females) | p-value = 0.7795 |
| **Panel 3C** | *Scn1a*$^{+/-}$DMSO | *Scn1a*$^{+/-}$ KU | Mann-Whitney test |
| Seizure severity score mean ± SEM | 5.7 ± 0.17 N=8 mice (4 males, 4 females) | 5.75 ± 0.16 N=8 mice (4 males, 4 females) | p-value = >0.9999 |
| **Panel 3D** | *Scn1a*$^{+/-}$-CTRL | *Scn1a*$^{+/-}$-BUM | Mann-Whitney test |
| Threshold temperature (°C) mean ± SEM | 41.79 ± 0.24 N = 9 mice (5 males, 4 females) | 41.81 ± 0.17 N = 9 mice (4 males, 5 females) | p-value = >0.9999 |
| **Panel 3E** | *Scn1a*$^{+/-}$-CTRL | *Scn1a*$^{+/-}$-BUM | Mann-Whitney test |
| Seizure severity score mean ± SEM | 6 ± 0 N = 9 mice (5 males, 4 females) | 6 ± 0 N = 9 mice (4 males, 5 females) | p-value = >0.9999 |
| **Panel 3F** | *Scn1a*$^{+/-}$-DMSO | *Scn1a*$^{+/-}$-KU | Mann-Whitney test |
| Seizure frequency (seizures/day) mean ± SEM | 0.91 ± 024 N=11 mice (4 males, 7 females) | 0.66±0.18 N=11 mice (4 males, 7 females) | p-value = 0.6633 |
| **Panel 3G** | *Scn1a*$^{+/-}$-DMSO | *Scn1a*$^{+/-}$-KU | Mann-Whitney test |
| Clusters of Seizures (number) mean ± SEM | 1.18±0.35 (4 males, 7 females) | 1.09 ± 0.41 (4 males, 7 females) | p-value = 0.7575 |
| **Panel 3I** | *Scn1a*$^{+/-}$-CTRL | *Scn1a*$^{+/-}$- BUM | Mann-Whitney test |
| Seizure frequency (seizures/day) mean ± SEM | 0.48± 0.10 N=12 mice (7 males, 5 females) | 0.61±0.18 N=13 mice (8 males, 5 females) | p-value = 0.8831 |
| **Panel 3J** | *Scn1a*$^{+/-}$-CTRL | *Scn1a*$^{+/-}$- BUM | Mann-Whitney test |
| Clusters of Seizures (number) mean ± SEM | 0.83 ± 0.30 N=12 mice (7 males, 5 females) | 0.54 ± 0.24 N=13 mice (8 males, 5 females) | p-value = 0.5005 |

#### Figure 4

| **Panel 4B** | WT-DMSO | *Scn1a$^{+/-}$*-DMSO | Scn1a$^{+/-}$-KU | One-Way ANOVA, Fisher's LSD post-hoc test |
|---|---|---|---|---|
| sociability index males mean ± SEM | 0.771±0.044 N=11 mice | -0.069±0.165 N=7 mice | 0.623±109 N=6 mice | p-value: WT-DMSO vs Scn1a$^{+/-}$-DMSO = **<0.0001**, WT-DMSO vs. Scn1a$^{+/-}$-KU = 0.3208, Scn1a$^{+/-}$-DMSO vs. Scn1a$^{+/-}$-KU = **0.0003** |
| **Panel 4C** | WT-DMSO | Scn1a$^{+/-}$-DMSO | Scn1a$^{+/-}$-KU | One-Way ANOVA, Fisher's LSD post-hoc test |
| sociability index females mean ± SEM | 0.574±0.062 N=11 mice | 0.489 ± 0.126 N=7 mice | 0.556±0.110 N=8 mice | p-value: WT-DMSO vs Scn1a$^{+/-}$-DMSO = 0.5269, WT-DMSO vs. Scn1a$^{+/-}$-KU = 0.8881, Scn1a$^{+/-}$-DMSO vs. Scn1a$^{+/-}$-KU = 0.6410 |
| **Panel 4d** | WT-CTRL | *Scn1a$^{+/-}$*-CTRL | *Scn1a$^{+/-}$*-BUM | One-Way ANOVA, Fisher's LSD post-hoc test |
| sociability index males mean ± SEM | 0.610±0.082 N=11 mice | 0.182±0.079, N=9 mice | 0.488±0.055 N=13 mice | p-value: WT-CTRL vs Scn1a$^{+/-}$-CTRL = **0.0009**, WT-CTRL vs. Scn1a$^{+/-}$-BUM = 0.2046, Scn1a$^{+/-}$-CTRL vs. Scn1a$^{+/-}$-BUM = **0.0252** |

#### Figure 5

| **Panel 5A** | WT-DMSO | Scn1a$^{+/-}$-DMSO | Scn1a$^{+/-}$-KU | One-Way ANOVA, Fisher's LSD post-hoc test |
|---|---|---|---|---|
| % alternation mean ± SEM | 49.03 ± 2.98, N = 14 | 36.88 ± 4.79, N = 10 | 38.58 ± 4.18, N = 9 | p-value: WT-DMSO vs Scn1a$^{+/-}$-DMSO = **0.0295**, WT-DMSO vs. Scn1a$^{+/-}$-KU = 0.0664, Scn1a$^{+/-}$-DMSO vs. Scn1a$^{+/-}$-KU = 0.7748 |
| **Panel 5B** | WT-DMSO | Scn1a$^{+/-}$-DMSO | Scn1a$^{+/-}$-KU | One-Way ANOVA, Fisher's LSD post-hoc test |
| n° entries mean ± SEM | 30.57 ± 2.46, N = 14 | 48.80 ± 7.13, N = 10 | 45.44 ± 2.71, N = 9 | p-value: WT-DMSO vs Scn1a+/--DMSO = **0.0046**, WT-DMSO vs. Scn1a+/--KU = **0.0218**, Scn1a+/--DMSO vs. Scn1a+/--KU = 0.6154 |
| **Panel 5C** | WT-DMSO | Scn1a$^{+/-}$-DMSO | Scn1a$^{+/-}$-KU | One-Way ANOVA, Fisher's LSD post-hoc test |
| Total distance mean ± SEM | 14.85 ± 1.14, N = 14 | 24.83 ± 3.98, N = 10 | 20.95 ± 0.90, N = 9 | p-value: WT-DMSO vs Scn1a$^{+/-}$-DMSO = **0.0034**, WT-DMSO vs. Scn1a$^{+/-}$-KU = **0.0694**, Scn1a$^{+/-}$-DMSO vs. Scn1a$^{+/-}$-KU = 0.2741 |
| **Panel 5D** | WT-CTRL | Scn1a$^{+/-}$-CTRL | Scn1a$^{+/-}$-BUM | One-Way ANOVA, Fisher's LSD post-hoc test |

| | | | | |
|---|---|---|---|---|
| % alternation mean ± SEM | 50.81 ± 4.55, N=10 | 29.48 ± 3.24, N=9 | 41.96 ± 2.60, N=13 | p-value: WT-CTRL vs Scn1a$^{+/-}$-CTRL = **0.0003**, WT-CTRL vs. Scn1a$^{+/-}$-BUM = 0.0719, Scn1a$^{+/-}$-CTRL vs. Scn1a$^{+/-}$-BUM = **0.0160** |
| **Panel 5E** | WT-CTRL | Scn1a$^{+/-}$-CTRL | Scn1a$^{+/-}$-BUM | One-Way ANOVA, Fisher's LSD post-hoc test |
| n°entries mean ± SEM | 27.90 ± 2.795, N = 10 | 51.67 ± 5.069, N = 9 | 36.85 ± 3.596, N = 13 | p-value: WT-CTRL vs Scn1a$^{+/-}$-CTRL = **0.0003**, WT-CTRL vs. Scn1a$^{+/-}$-BUM = 0.1009, Scn1a$^{+/-}$-CTRL vs. Scn1a$^{+/-}$-BUM = **0.0109** |
| **Panel 5F** | WT-CTRL | Scn1a$^{+/-}$-CTRL | Scn1a$^{+/-}$-BUM | One-Way ANOVA, Fisher's LSD post-hoc test |
| Total distance mean ± SEM | 13.77 ± 1.55, N = 10 | 23.85 ± 1.90, N = 9 | 17.14 ± 1.68, N = 13 | p-value: WT-CTRL vs Scn1a$^{+/-}$-CTRL = **0.0005**, WT-CTRL vs. Scn1a$^{+/-}$-BUM = 0.1646, Scn1a$^{+/-}$-CTRL vs. Scn1a$^{+/-}$-BUM = **0.0100** |
| **Panel 5G** | WT-DMSO | Scn1a$^{+/-}$-DMSO | Scn1a$^{+/-}$-KU | One-Way ANOVA, Fisher's LSD post-hoc test |
| % alternation mean ± SEM | 47.92 ± 3.41, N = 16 | 42.14 ± 4.09, N = 7 | 43.66 ± 6.55, N = 9 | p-value: WT-DMSO vs Scn1a$^{+/-}$-DMSO = 0.4047, WT-DMSO vs. Scn1a$^{+/-}$-KU = 0.5035, Scn1a$^{+/-}$-DMSO vs. Scn1a$^{+/-}$-KU = 0.8426 |
| **Panel 5H** | WT-DMSO | Scn1a$^{+/-}$-DMSO | Scn1a$^{+/-}$-KU | One-Way ANOVA, Fisher's LSD post-hoc test |
| n°entries mean ± SEM | 27.63 ± 1.52, N = 16 | 52.14 ± 9.50, N = 7 | 37.11 ± 5.48, N = 9 | p-value: WT-DMSO vs Scn1a$^{+/-}$-DMSO = **0.0011**, WT-DMSO vs. Scn1a$^{+/-}$-KU = 0.1394, Scn1a$^{+/-}$-DMSO vs. Scn1a$^{+/-}$-KU = 0.0560 |
| **Panel 5I** | WT-DMSO | Scn1a$^{+/-}$-DMSO | Scn1a$^{+/-}$-KU | One-Way ANOVA, Fisher's LSD post-hoc test |
| Total distance mean ± SEM | 13.50 ± 0.61, N = 16 | 26.06 ± 4.94, N = 7 | 19.11 ± 2.33, N = 9 | p-value: WT-DMSO vs Scn1a$^{+/-}$-DMSO = **0.0006**, WT-DMSO vs. Scn1a$^{+/-}$-KU = 0.0718, Scn1a$^{+/-}$-DMSO vs. Scn1a$^{+/-}$-KU = 0.0657 |
| **Panel 5J** | WT-CTRL | Scn1a$^{+/-}$-CTRL | Scn1a$^{+/-}$-BUM | One-Way ANOVA, Fisher's LSD post-hoc test |
| % alternation mean ± SEM | 53.98 ± 7.04, N = 9 | 43.23 ± 3.15, N = 8 | 42.75 ± 3.60, N = 10 | p-value: WT-CTRL vs Scn1a$^{+/-}$-CTRL = 0.1523, WT-CTRL vs. Scn1a$^{+/-}$-BUM = 0.0808, Scn1a$^{+/-}$-CTRL vs. Scn1a$^{+/-}$-BUM = 0.8037 |
| **Panel 5K** | WT-CTRL | Scn1a$^{+/-}$-CTRL | Scn1a$^{+/-}$-BUM | One-Way ANOVA, Fisher's LSD post-hoc test |

| | | | | |
|---|---|---|---|---|
| N°entries mean ± SEM | 22.44 ± 3.09, N = 9 | 51.13 ± 8.60, N = 8 | 36.09 ± 3.85, N = 10 | p-value: WT-CTRL vs Scn1a$^{+/-}$-CTRL = **0.0008**, WT-CTRL vs. Scn1a$^{+/-}$-BUM = 0.1335, Scn1a$^{+/-}$-CTRL vs. Scn1a$^{+/-}$-BUM = **0.0228** |
| **Panel 5L** | WT-CTRL | Scn1a$^{+/-}$-CTRL | Scn1a$^{+/-}$-BUM | One-Way ANOVA, Fisher's LSD post-hoc test |
| Total distance mean ± SEM | 10.61 ± 1.28, N = 9 | 23.34 ± 3.05, N = 8 | 17.70 ± 1.67, N = 10 | p-value: WT-CTRL vs Scn1a$^{+/-}$-CTRL = **0.0001**, WT-CTRL vs. Scn1a$^{+/-}$-BUM = **0.0346**, Scn1a$^{+/-}$-CTRL vs. Scn1a$^{+/-}$-BUM = **0.0165**) |

**Figure 6**

| | | | | | |
|---|---|---|---|---|---|
| **Panel 6A** | WT P7 | Scn1a$^{+/-}$ P7 | WT P10-11 | Scn1a$^{+/-}$ P10-11 | Mann-Whitney test |
| Righting reflex latency mean ± SEM | 8.98 ± 2.65, N = 16 (6 males, 10 females) | 4.73 ± 1.91, N = 15 (6 males, 9 females) | 1.37 ± 0.17, N = 12 (5 males, 7 females) | 1.45 ± 0.30, N = 13 (6 males, 7 females) | p-value: P7 = 0.0723, P10-11 >0.9999 |
| **Panel 6B** | WT P7 | Scn1a$^{+/-}$ P7 | WT P10-11 | Scn1a$^{+/-}$ P10-11 | Fisher's exact test |
| Grasping reflex | N = 12 (6 males, 6 females) | N = 13 (6 males, 7 females) | N = 12 (6 males, 6 females) | N = 13 (6 males, 7 females) | p-value: P7 = >0.9999, P10-11 >0.9999 |
| **Panel 6C** | WT | Scn1a$^{+/-}$ | Mann-Whitney test | | |
| Negative geotaxis time mean ± SEM | 16.84 ± 3.66, N=12 (6 males, 6 females) | 15.80 ± 4.08, N = 13 (6 males, 7 females) | p-value = 0.3435 | | |
| **Panel 6D** | WT | Scn1a$^{+/-}$ | Mann-Whitney test | | |
| Gait latency mean ± SEM | 22.89 ± 2.10, N = 12 (6 males, 6 females) | 22.65 ± 2.49, N = 13 (6 males, 7 females) | p-value = 0.7391 | | |
| **Panel 6E** | WT | Scn1a$^{+/-}$ | Fisher's exact test | | |
| Olfactory motivation %success | N = 16 (9 males, 7 females) | N = 19 (9 males, 10 females) | p-value = **0.0184** | | |
| Time to reach the | 36.70 ± 2.853 | 41.89 ± 2.184 | Mann-Whitney test<br>p-value = 0.1846 | | |

| | | | |
|---|---|---|---|
| nest mean ± SEM | N = 16 (9 males, 7 females) | N = 19 (9 males, 10 females) | |
| **Panel 6F** | WT | Scn1a$^{+/-}$ | Fisher's exact test |
| Eye opening | N = 16 (5 males, 11 females) | N = 17 (7 males, 10 females) | p-value = **0.0457** |

**Figure 7**

| **Panel 7B** | WT | Scn1a$^{+/-}$ | unpaired t-test |
|---|---|---|---|
| sIPSC frequency (Hz) mean ± SEM | 2.219 ± 0.2165, n = 13 cells, N = 3 animals (2 males, 1 female) | 1.605 ± 0.1673, n = 18 cells, N = 3 animals (1 male, 2 females) | p-value = **0.0301** |
| **Panel 7C** | WT | Scn1a$^{+/-}$ | unpaired t-test |
| sEPSC frequency (Hz) mean ± SEM | 3.63 ± 0.39, n=15 cells, N = 3 animals (2 males, 1 female) | 3.15 ± 0.34, n = 17 cells, N = 3 animals (1 male, 2 females) | p-value = 0.3591 |
| **Panel 7D** | WT | Scn1a$^{+/-}$ | unpaired t-test |
| E/I ratio mean ± SEM | 1.398 ± 0.1957, n = 10 cells, N = 3 animals (2 males, 1 female) | 2.251 ± 0.3375, n = 16 cells, N = 3 animals (1 male, 2 females) | p-value = **0.0741** |
| **Panel 7E** | WT | Scn1a$^{+/-}$ | unpaired t-test |
| sIPSC amplitude (pA) mean ± SEM | 24.80 ± 1.41, n = 13 cells, N = 3 animals (2 males, 1 female) | 20.77 ± 1.90, n = 18 cells, N = 3 animals (1 male, 2 females) | p-value = 0.1243 |
| **Panel 7F** | WT | Scn1a$^{+/-}$ | unpaired t-test |
| sEPSC amplitude (pA) mean ± SEM | 23.59 ± 1.68, n = 14 cells, N = 3 animals (2 males, 1 female) | 21.27 ± 1.66, n = 17 cells, N = 3 animals (1 male, 2 females) | p-value = 0.3400 |

**Figure 8**

| **Panel 8Ca** | WT | Scn1a$^{+/-}$ | Mann-Whitney test |
|---|---|---|---|
| Spine density mean ± SEM | 0.5546 ± 0.02270; N = 9 mice (4 males, 5 females) | 0.6120 ± 0.01862; N mice = 9 (4 males, 5 females) | p-value = **0.0767** |
| **Panel 8Cb** | WT | Scn1a$^{+/-}$ | Mann-Whitney test |

| | | | |
|---|---|---|---|
| Spine head width<br>mean ± SEM | 0.5173 ± 0.03001; N = 8 mice (4 males, 4 females) | 0.3787 ± 0.01432; N = 9 mice (4 males, 5 females) | p-value = **0.0006** |
| **Panel 8Da** | WT | $Scn1a^{+/-}$ | Mann-Whitney test |
| Spine density<br>mean ± SEM | 0.9855 ± 0.02019; N = 10 mice (5 males, 5 females) | 0.9748 ± 0.02860; N =11 mice<br>(6 males, 5 females) | p-value = 0.9017 |
| **Panel 8Db** | WT | $Scn1a^{+/-}$ | Mann-Whitney test |
| Spine head width<br>mean ± SEM | 0.5084 ± 0.01938; N = 9 mice (4 males, 5 females) | 0.3944 ± 0.01959; N = 11 mice<br>(6 males, 5 females) | p-value = **0.0016** |
